# Nitrate-driven anaerobic oxidation of ethane and butane by bacteria

**DOI:** 10.1101/2023.08.24.554723

**Authors:** Mengxiong Wu, Jie Li, Chun-Yu Lai, Andy O Leu, Shengjie Sun, Rui Gu, Dirk V Erler, Lian Liu, Lin Li, Gene W. Tyson, Zhiguo Yuan, Simon J. McIlroy, Jianhua Guo

## Abstract

The short-chain gaseous alkanes (ethane, propane and butane; SCGAs) are important components of natural gas, yet our understanding of their fate in environmental systems is poorly understood. Microbially mediated anaerobic oxidation of SCGAs coupled to nitrate reduction has been demonstrated for propane, but is yet to be shown for ethane or butane – despite being energetically feasible. Here we report two independent bacterial enrichments performing anaerobic ethane and butane oxidation, respectively, coupled to nitrate reduction to dinitrogen gas and ammonium. Isotopic ^13^C-and ^15^N-labelling experiments, mass and electron balance tests, and metabolite and meta-omics analyses collectively reveal that the recently described propane-oxidising ‘*Candidatus* Alkanivorans nitratireducens’ was also responsible for nitrate-dependent anaerobic oxidation of the SCGAs in both these enrichments. The complete genome of this species encodes alkylsuccinate synthase genes for the activation of ethane/butane via fumarate addition. Further substrate range tests confirm ‘*Ca.* A. nitratireducens’ is metabolically versatile, being able to degrade ethane, propane and butane under anaerobic conditions. Moreover, our study proves nitrate as an additional electron sink for ethane and butane in anaerobic environments, and for the first time demonstrates the use of the fumarate addition pathway in anaerobic ethane oxidation. These findings significantly contribute to our understanding of microbial metabolism of SCGAs in anaerobic environments.

---

Short-chain gaseous alkanes (SCGAs), including ethane, propane and butane, are abundant components of natural gas (up to 20%) and contribute significantly to the formation of tropospheric ozone and secondary organic aerosols, ^1–3^, thus negatively impacting air quality and climate^4,5^. The atmospheric SCGA emissions have greatly increased since preindustrial times, reaching ∼10 Tg yr^-1^ for ethane, propane, butane and ∼4 Tg yr^-1^ for *iso*-butane^6,7^. Fortunately, microorganisms can utilize the SCGAs under aerobic and anaerobic conditions, significantly reducing their flux from natural ecosystems to the atmosphere^8,9^.

While the microbiology of aerobic oxidation of SCGAs has been well studied^10^, the microorganisms and metabolic pathways involved in the anaerobic oxidation of these gases have only been identified in recent years. The archaeal species ‘*Candidatus* Argoarchaeum ethanivorans’ and ‘*Candidatus* Syntrophoarchaeum’ oxidize ethane and butane via the formation of ethyl-or butyl-coenzyme M, respectively, in syntrophic consortia with sulfate-reducing bacteria (SRB) ^11,12^. In contrast, the deltaproteobacterial isolate *Desulfosarcina aeriophaga* BuS5 oxidize propane and butane via reaction with fumarate, generating propyl-and butyl-succinates (the fumarate addition pathway), coupled to the direct reduction of sulfate to sulfide^13,14^. Our recent study described a bacterial species ‘*Candidatus* Alkanivorans nitratireducens’ belonging to the Class of Symbiobacteriia that can oxidize propane via the fumarate addition pathway coupled to the reduction of nitrate to nitrite^15^. The oxidation of ethane and butane coupled to nitrate reduction is yet to be shown, but would also be thermodynamically feasible (Equations 1 and 2) and potentially important given the prevalence of nitrate in natural environments^16,17^.

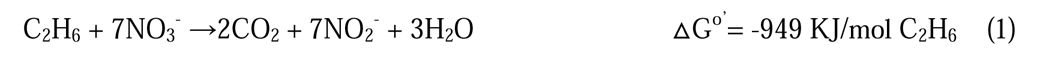

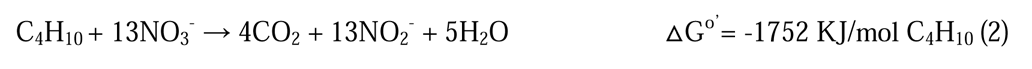

Especially, anaerobic ethane oxidation remains poorly understood, with direct evidence for this metabolic process limited to archaea^11,18^. Indeed, ethane activation mediated by bacteria has not been proven, in clear contrast to the multiple discoveries of SRB-mediated anaerobic propane and butane degradation^13,19,20^. The fumarate addition pathway is considered the most common mechanism for anaerobic degradation of hydrocarbons including propane, butane and various other n-alkanes ranging from C_6_ (n-hexane) to C_16_ (n-hexadecane) ^21–24^. The oxidation of ethane via this mechanism is also likely to occur in the environment, given ethyl-succinate, the signature metabolite generated by ethane activation via reaction with fumarate, is frequently detected in hydrocarbon-rich environments, such as crude oil production wells, coal beds and oilfields^25–27^. However, physiological evidence for anaerobic ethane oxidation via the fumarate addition pathway is lacking.

In this study, we address these knowledge gaps by enriching microbial consortia able to couple anaerobic ethane and butane oxidation to nitrate reduction, and characterizing the key metabolic pathways via a multi-omics approach (metagenomics, metatranscriptomics and metaproteomics). The alkane oxidising population in both enrichments is the same species as anaerobic propane degrading strain ‘*Ca.* A. nitratireducens’ identified previously^15^, and is suggested to mediate ethane and butane oxidation via reactions with fumarate.

## Enrichment cultures able to mediate nitrate-dependent anaerobic oxidation of ethane and butane

In this study, two anaerobic bioreactors seeded with activated sludge and anaerobic digestion sludge from a wastewater treatment plant were operated for more than 1,000 days. One was fed with ethane (C_2_H_6_) and nitrate, whilst the other with butane (C_4_H_10_) and nitrate. The C_2_H_6_-fed bioreactor showed simultaneous consumption of C_2_H_6_ and nitrate, with production of dinitrogen gas and ammonium, and transient accumulation of nitrite (Supplementary Fig. 1a). Similarly, nitrate consumption and ammonium production were observed in the C_4_H_10_-fed reactor (Supplementary Fig. 1b). No nitrate consumption was observed in the control incubations without the addition of either C_2_H_6_ or C_4_H_10_ or enrichment culture biomass (Supplementary Fig. 2), indicating that nitrate reduction (to nitrite and ammonium) was a biological process and coupled to the consumption of these alkanes.

Stoichiometric experiments were conducted directly in the parent C_2_H_6_-fed reactor or with subcultures from the parent C_4_H_10_-fed reactor to establish nitrogen and electron balances. The reduction of NO_3_^-^ proceeded in two distinct phases for both C_2_H_6_-and C_4_H_10_-fed systems (Fig. 1a, 1b, Supplementary Fig. 3). In Phase 1, NO_3_^-^ was reduced to NO_2_^-^ and N_2_ with negligible NH_4_^+^ accumulation (Equations 1, 2, 4 and 5). In Phase 2, when NO_3_^-^ was depleted, NO_2_^-^ was further reduced to NH_4_^+^ and N_2_ (Equations 2, 3, 5 and 6). The total amounts of the produced nitrogen species (NH_4_^+^ + N_2_) for C_2_H_6_-(1.93 ± 0.15 mmol N/L) and C_4_H_10_-fed (1.53 ± 0.08 mmol N/L) batch tests were close to the amounts of nitrogen oxyanions consumed (NO_3_^-^ + NO_2_^-^, 1.77 ± 0.05 and 1.63 ± 0.10 mmol N/L for C_2_H_6_ and C_4_H_10_-fed cultures, respectively, Fig. 1c, 1d, Supplementary Table 1). This indicates NH_4_^+^ and N_2_ were the final products generated from NO_3_^-^ and NO_2_^-^ reduction. The amounts of electrons required for denitrification (NO_3_^-^ reduction to N_2_) and dissimilatory nitrate reduction to ammonia (DNRA) in the C_2_H_6_-and C_4_H_10_-fed batch tests represent 92 ± 4% and 99 ± 6% of the maximum electrons available in C_2_H_6_ and C_4_H_10_ oxidation to CO_2_, respectively (Fig. 1c, 1d, Supplementary Table 1), suggesting electrons were mainly diverted to NO_3_^-^ reduction in these systems.

**Fig. 1.**
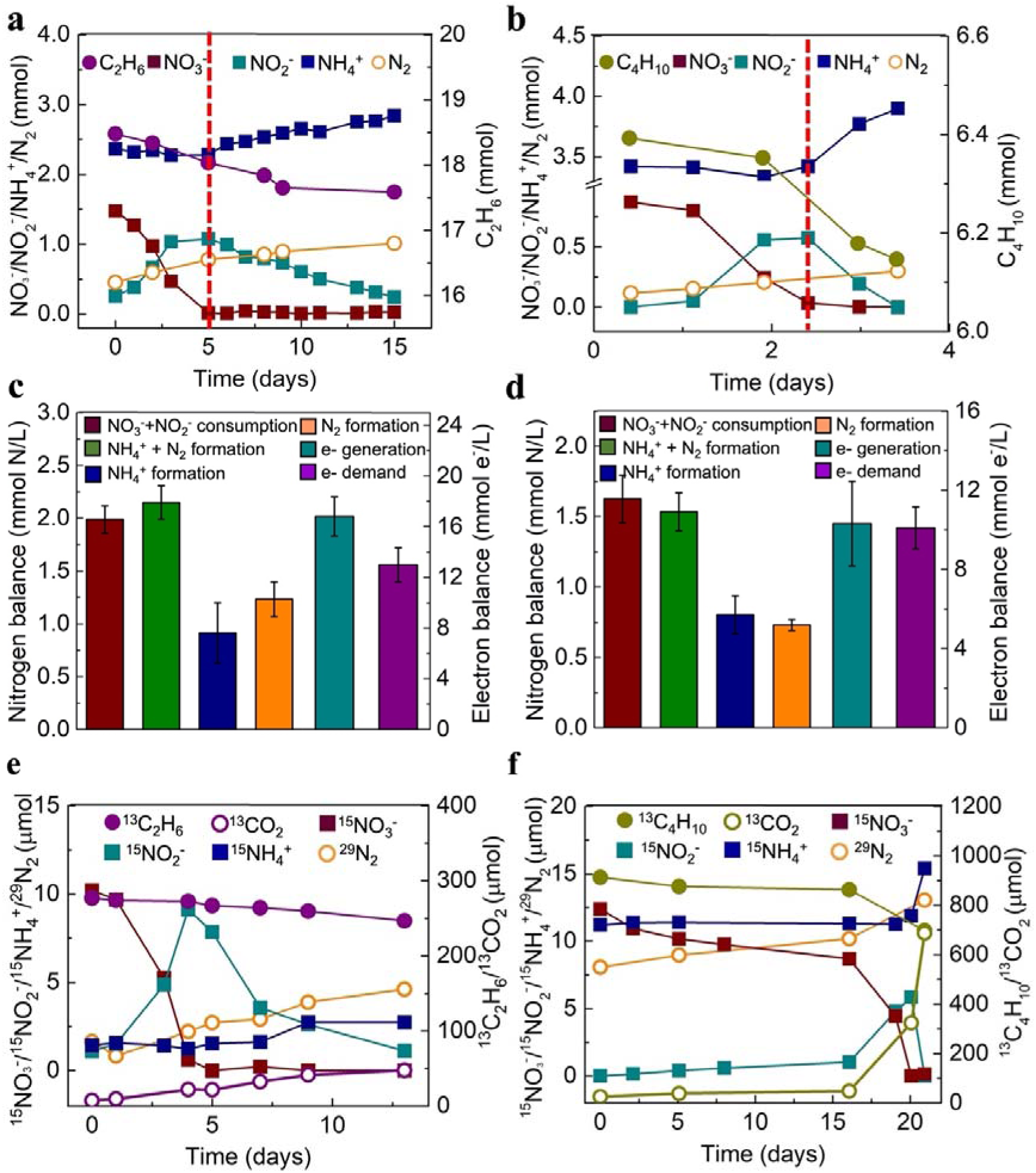
Mass and electron balance batch tests, and isotope labelling experiments confirmed anaerobic ethane/butane oxidation was coupled to nitrate reduction by the bioreactor enrichment cultures fed with C_2_H_6_/C_4_H_10_. **a, b,** Typical biochemical profiles of the ethane **(a,** started on Day 490) and butane (**b,** started on Day 1,100) systems showing simultaneous nitrate and ethane/butane consumption with transitory formation of nitrite, and production of dinitrogen gas and ammonium. There were two distinct phases for NO_3_^-^ reduction. In Phase 1, NO_3_^-^ was reduced to NO_2_^-^ and N_2_, with negligible NH_4_^+^ production; while in Phase 2, the accumulated NO ^-^ was reduced to both N and NH ^+^. **c, d,** Average nitrogen-and electron balances calculated from the three batch tests for C_2_H_6_- (c) and C_4_H_10_-**(d)** fed bioreactors (Supplementary Table 1 shows the complete data and calculation). Error bars represent standard errors from biological triplicates. **e, f,** oxidation of ^13^C_2_H_6_ **(e)** or ^13^C_4_H_10_ **(f)** to ^13^CO_2_, and reduction of ^15^NO_3_^-^ to ^15^NH_4_^+^ and ^29^N_2_ with temporary generation of ^15^NO_2_^-^ during the isotope labelling test.

To verify the final products of anaerobic C_2_H_6_/C_4_H_10_ oxidation coupled to nitrate reduction, subcultures from the parent reactors were incubated with ^13^C-labelled C_2_H_6_ (^13^CH_3_^13^CH_3_) or C_4_H_10_ (^13^CH_3_^13^CH_2_^13^CH_2_^13^CH_3_) and ^15^N-labelled nitrate (^15^NO_3_^-^) in 0.6 L glass vessels. Concomitant to ^13^C_2_H_6_ /^13^C_4_H_10_ consumption, ^13^CO_2_ was produced in both tests. The amounts of ^13^CO_2_ produced from the labelled C_2_H_6_- (40 µmol) and C_4_H_10_-fed (661 µmol) batches were 67% and 77%, respectively, of the consumed ^13^C in ^13^C_2_H_6_ (60 µmol) and ^13^C_4_H_10_ (840 µmol) (Fig. 1e, 1f). Similarly, the total amounts of CO_2_ produced were 71% and 83% of total consumed carbon in C_2_H_6_ and C_4_H_10_, respectively (Supplementary Fig. 4). These results suggest CO_2_ was the dominant end product from C_2_H_6_ and C_4_H_10_ oxidation. The total ^15^N in ^29^N_2_, ^30^N_2_, and ^15^NH_4_^+^ produced (8.9 and 9.2 µmol in total in the C_2_H_6_ and C_4_H_10_-fed batch, respectively) was concordant with the totally consumed ^15^NO_3_^-^ (10.2 and 12.2 µmol for C_2_H_6_ and C_4_H_10_-fed batches, respectively), confirming the reduction of NO_3_^-^ to N_2_ and NH_4_^+^ (Fig. 1e, 1f). These findings collectively support nitrate-dependent anaerobic oxidation of C_2_H_6_ and C_4_H_10_ in the two bioreactors (Equations 1-6).

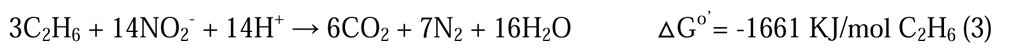

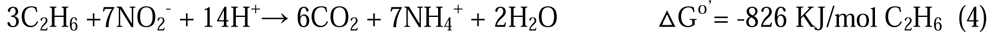

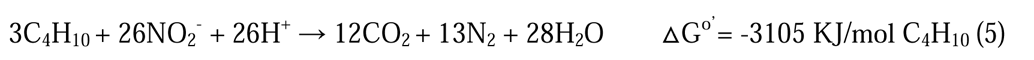

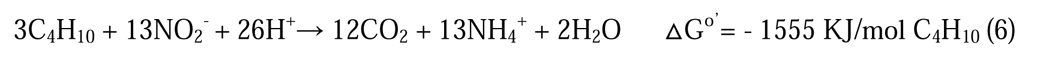

### Microbial community structure and genome recovery

16S rRNA gene amplicon sequencing of the biomass from both bioreactor enrichments revealed the dominance of the recently described propane oxidizing firmicute ‘*Ca.* A. nitratireducens’^15^ in both systems (100% amplicon sequence similarity; 5.3-10.5% abundance for C_2_H_6_-fed reactor and 4.3-18.3% for C_4_H_10_-fed reactor, Supplementary Fig. 5). The metagenomes of both cultures were obtained by applying both long (Nanopore) and short read (Illumina) sequencing for biomass samples collected from the C_2_H_6_- (on Day 746) and C_4_H_10_-fed (Day 1,150) bioreactors (Supplementary Table 2). In total, 63 and 37 high-quality genomes (≥70% completeness and ≤10% contamination based on CheckM) were retrieved for the C_2_H_6_- and C_4_H_10_-fed bioreactor enrichments, respectively (Supplementary Data 1). These included two complete circularised genomes of the dominant ‘*Ca*. A. nitratireducens’ in the C_2_H_6_- (15.0% of relative abundance, a size of 2.42 Mbp, Supplementary Table 3, Supplementary Fig. 6) and C_4_H_10_-fed (16.7% of relative abundance, a size of 2.32 Mbp, Supplementary Table 4, Supplementary Fig. 6) bioreactors. These genomes had average nucleotide identities (ANI) of 99.96% and 99.55%, and average amino acid identities (AAI) of 99.96% and 99.57% (Supplementary Fig. 7) to the ‘*Ca.* A. nitratireducens’ genome previously recovered from the C_3_H_8_-fed culture^15^, confirming that the three genomes likely represent the same species^28^.

### Analyses of metabolic pathways of ‘*Ca*. A. nitratireducens’

Consistent with ‘*Ca.* A. nitratireducens’ originating from the C_3_H_8_-fed system (referred to as Strain P), the closed genomes of ‘*Ca*. A. nitratireducens’ in C_2_H_6_- and C_4_H_10_-fed bioreactors (referred to as Strains E and B) both contain three alkylsuccinate synthase catalytic subunits (AssA, Supplementary Fig. 8a) which are phylogenetically distant from other available fumarate addition enzymes in the UniProt database (Supplementary Fig. 8b). To support the role of these alkylsuccinate synthase complexes in ethane/butane oxidation, key metabolites from the active cultures were analysed by ultra-high-sensitivity triple quadrupole mass spectrometry. A mass peak (m/z: 275>73.1) at the retention time of 9.940 min was detected for the C_2_H_6_-fed bioreactor, corresponding to the ethyl-succinate standard (Fig. 2a). Also, a mass peak (m/z: 303.0>147.1) at the retention time of 12.245 min was detected for the C_4_H_10_- fed bioreactor, corresponding to the butyl-succinate standard (Fig. 2b). These findings support that ethane/butane were activated by addition of fumarate, thus generating ethyl/butyl-succinate, which is consistent with the action of the alkylsuccinate synthase.

**Fig. 2.**
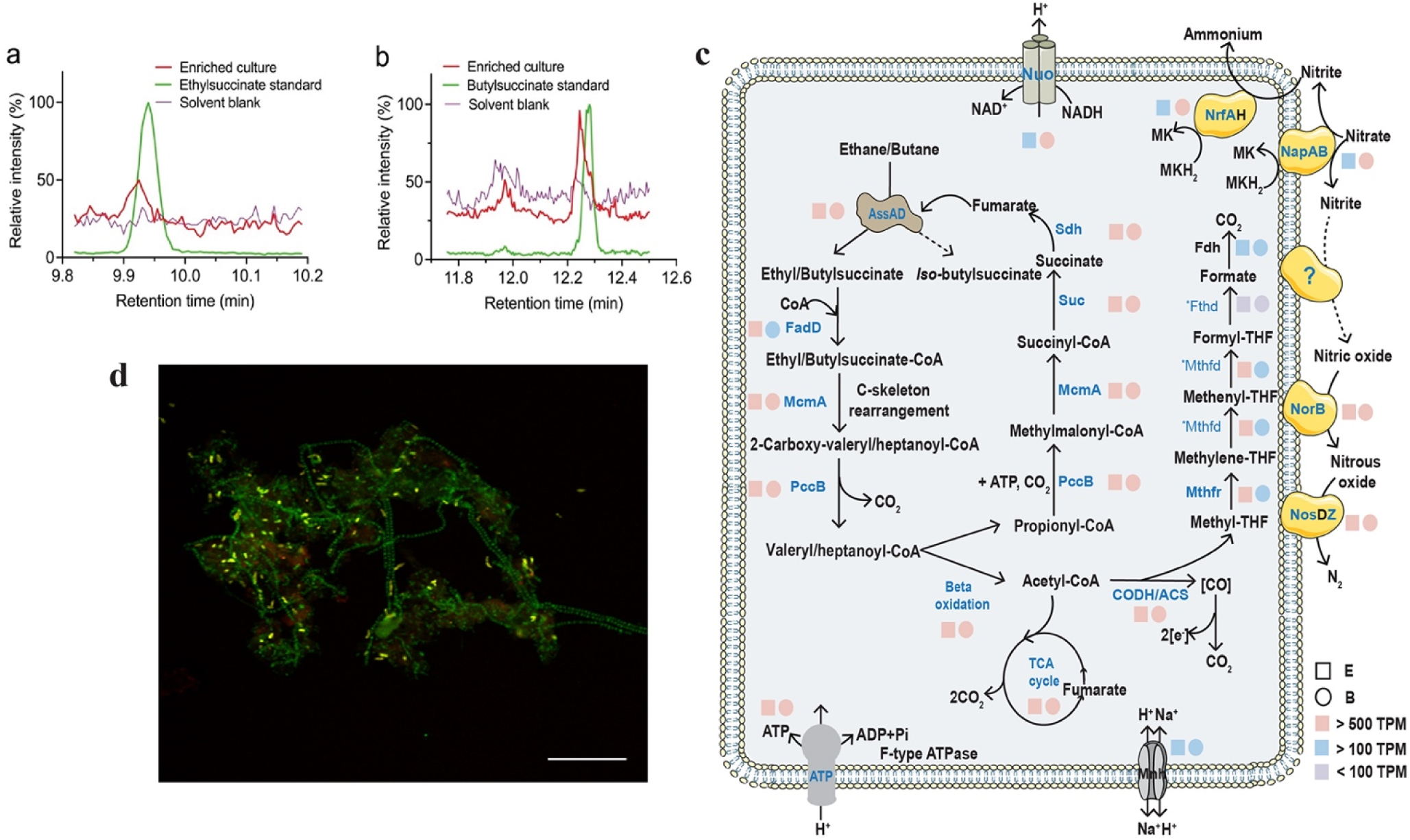
Metabolic intermediates, inferred metabolic pathways and a fluorescent micrograph of ‘*Ca.* A. nitratireducens’. **a,** Partial ion chromatograms (ion transition, m/z: 275>73.1) of culture extracts from the C_2_H_6_-fed bioreactor displayed a characteristic peak at a retention time of 9.940 min, matching the ethylsuccinate standard. **b,** A characteristic peak at a retention time of 12.245 min (ion transition, m/z: 303.0>147.1), consistent with the peak from the butyl-succinate standard, was observed for the culture extracts from the C_4_H_10_-fed bioreactor (n=4 at different sampling points). **c,** Cell cartoon illustrating ‘*Ca.* A. nitratireducens’ in the C_2_H_6_- or C_4_H_10_-fed bioreactors (E or B) use alkylsuccinate synthase to activate ethane/butane to ethyl/butyl-succinate, which are further converted to acetyl-CoA and propionyl-CoA. Fumarate could be regenerated by the methylmalonyl-CoA pathway or the tricarboxylic acid (TCA) cycle. CO_2_ is produced through the TCA cycle or the reverse Wood–Ljungdahl pathway. The E and B both harbour genes that enable denitrification (except *nirS/K*) and dissimilatory nitrate reduction to ammonium. The colour of the square and circle symbols indicates the normalized gene expression values calculated as TPM (total transcripts per million). Blue bold text shows that the proteins were fully or partially detected in the protein extracts (*Mthfd and *Fthd were only identified in B and E, respectively), while proteins in black text were not detected. **d,** A composite fluorescence micrograph of the C_4_H_10_-fed enrichment culture hybridized with the SYMB-1018 probe^15^ (Cy3, red; targeting ‘*Ca.* A. nitratireducens’) and EUBmix probe set^35^ (Fluorescein isothiocyanate label, green; All bacteria). ‘*Ca.* A. nitratireducens’ cells appear yellow (red + green) and other bacterial cells appear green. The scale bar indicates 20Cμm. The representative image was selected based on the visual assessment of >3 separate hybridisation experiments. FISH was performed as detailed in our previous study^15^.

The genomes of Strains E and B also harbour other key genes involved in the further degradation of ethyl/butyl-succinate, including the methylmalonyl-CoA mutase genes (*mcmA*) for carbon-skeleton rearrangement, the propionyl-CoA carboxylase genes (*pccB*) for decarboxylation and the genes for beta-oxidation (Supplementary Data 2, 3, Fig. 2c). The propionyl-CoA generated from beta-oxidation could enter the methylmalonyl-CoA pathway to regenerate fumarate for subsequent rounds of ethane/butane activation. The acetyl-CoA may be completely oxidized to CO_2_ or used for fumarate regeneration via the oxidative tricarboxylic acid (TCA) cycle. CO_2_ can also be generated by oxidation of acetyl-CoA through the reverse Wood–Ljungdahl (WL) pathway for Strains E and B (Supplementary Data 2, 3, Fig. 2c), consistent with that proposed for Strain P and sulfate-dependent propane oxidizer—*Desulfosarcina aeriophaga* BuS5^14,15^. The metatranscriptomic and metaproteomic data indicated the E and B strains expressed the proposed fumarate addition pathway for complete ethane/butane oxidation to CO_2_ after alkane additions (Supplementary Data 2, 3, Fig. 2c). The ‘*Ca.* A. nitratireducens’ dominated the transcriptome profile of both the C_2_H_6_- (61.5% of the total transcriptome reads, Supplementary Table 5) and C_4_H_10_-fed bioreactors (84.5% the total transcriptome reads, Supplementary Table 6), indicating that they are the main drivers of anaerobic alkane oxidation in these systems.

Similar to Strain P, the genomes of Strains E and B both harbour and express genes encoding nitrate reductase (*napAB*) and cytochrome *c* nitrite reductases (*nrfAH*) required for DNRA process (Supplementary Data 2, 3, Fig. 2c). The expression of *nrfAH* was much higher in Strain B than E in Phase 2, consistent with the significantly higher DNRA rates (*p*<0.05) in the C_4_H_10_-fed bioreactor (0.77 ± 0.27 mmol/L/d) compared to the C_2_H_6_-fed bioreactor (0.11 ± 0.08 mmol/L/d). The NapAB and NrfA were also identified in protein extracts from both the C_2_H_6_- and C_4_H_10_-fed cultures (Supplementary Data 2, 3, Fig. 2c), further supporting Strains E and B were performing DNRA in these systems. The closed genomes of Strains E and B both lack nitric oxide-producing nitrite reductase (*nirS/K*), but encode nitric oxide reductase (*norB*) and nitrous oxide reductase (*nosZD*), consistent with Strain P. The *norB* and *nosD* genes were expressed and detected in the protein extracts for both E and B (Supplementary Data 2, 3, Fig. 2c), suggesting the active roles of these strains in the reduction of nitric oxide to dinitrogen gas. The phenomena that dinitrogen gas was generated without the apparent invovlement of *nirS/K* for the dominant ‘Ca. A. nitratireducens’ in all three systems, indicates that this species may indeed utilize a novel gene or novel pathway to reduce nitrite to nitric oxide^15^.

### SCGA metabolic versatility of ‘*Ca*. A. nitratireducens’

Structural modelling and molecular dynamics (MD) simulations were conducted to understand the potential functions of different AssAs in ‘*Ca.* A. nitratireducens’. The genome of Strain E encodes three AssAs that are 852aa in length with differing AAI between them (90.96-96.60%, Supplementary Table 7). The shorter AssA genes identified in ‘*Ca.* A. nitratireducens’ genomes of P and B were found to be due to open reading frame calling issues^29^ via full length alignments of the Strain E AssA genes to the AssA regions of the P and B MAGs, and manual identification of the start and stop codon. Further analyses of the AssAs in three strains show that full length alignment to the conserved domain (cd01677) for pyruvate formate lyase 2 and related enzymes is only found in the 852aa AssAs^30^, suggesting these AssAs are more likely to be complete. Given the overall high AssA gene similarities between strains (Supplementary Table 8), the three complete AssA genes in Strain E were used for structural modelling and MD simulations.

The MD results suggest AssA1 cannot stably bind to the key substrate—fumarate (movie S1), while AssA2 and AssA3 can form stable binding complexes with fumarate and ethane/propane/butane (movie S2-7, Fig. 3a-3f, Supplementary Fig. 9). Hydrogen bonding networks were found to be critical for the SCGA and fumarate bindings (Fig. 3a-3f, Supplementary Fig. 10). In addition, the putative radical sites Cys489 and Gly828 are situated at the core of AssA2/AssA3 and close to each other in all binding complexes (Fig. 3a-3f). These characteristics were suggested to be important for radical transfers in glycyl radical enzymes^31,32^, indicating that the radical transfer pathway may govern fumarate addition in AssA2/AssA3.

**Fig. 3.**
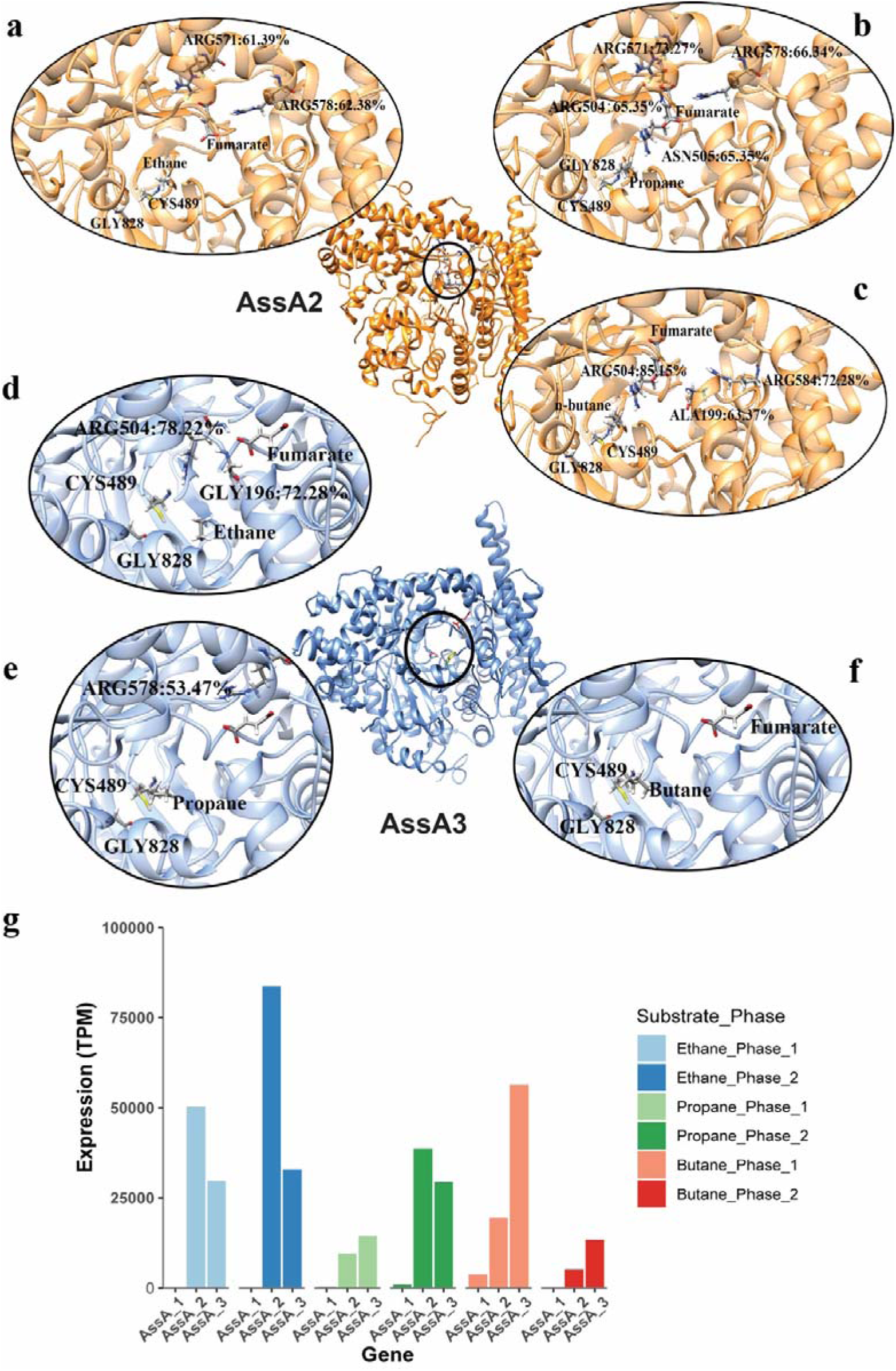
The molecular dynamics simulations and gene expression of alkylsuccinate synthases (AssAs) in ‘*Ca.* A. nitratireducens’. **a-f,** The structural representations of binding complexes of AssA2 with fumarate and ethane (**a**)/propane (**b**)/butane (**c**), and AssA3 with fumarate and ethane (**d**)/propane (**e**)/butane (**f**). Key residues of Cys489 and Gly828 are close to each other in all systems. Residues with occupancy of hydrogen bonds > 50% were also included in the figures. **g,** The normalized gene expression values of assA genes in ‘*Ca.* A. nitratireducens’ from C_2_H_6_-, C_3_H_8_- and C_4_H_10_-fed systems (calculated as total transcripts per million).

Metatranscriptomic profiles of the ethane, propane, and butane systems were mapped onto Strain E MAG to ensure consistency of the AssA gene lengths. In support of the MD results, AssA1 are relatively lowly expressed in all systems (Fig. 3g). However, the expression levels of AssA2 and AssA3 are relatively high in all systems (Fig. 3g), suggesting these proteins are more likely responsible for SCGA activation by ‘*Ca.* A. nitratireducens’.

To further validate if ‘Ca. A. nitratireducens’ is indeed able to oxidize all three SCGAs, substrate range tests were conducted for the C_2_H_6_-, C_3_H_8_- and C_4_H_10_-fed cultures. Incubation of subcultures from the C_2_H_6_-, C_3_H_8_- and C_4_H_10_-fed bioreactors with the other two SCGAs showed obvious ethane/propane/butane oxidation coupled to nitrate reduction to dinitrogen gas and ammonium (Fig. 4a-4f, Supplementary Fig. 11). These results provide compelling evidence that ‘*Ca*. A. nitratireducens’ has the metabolic versatility to oxidize the three tested SCGAs using nitrate as a terminal electron acceptor.

**Fig. 4.**
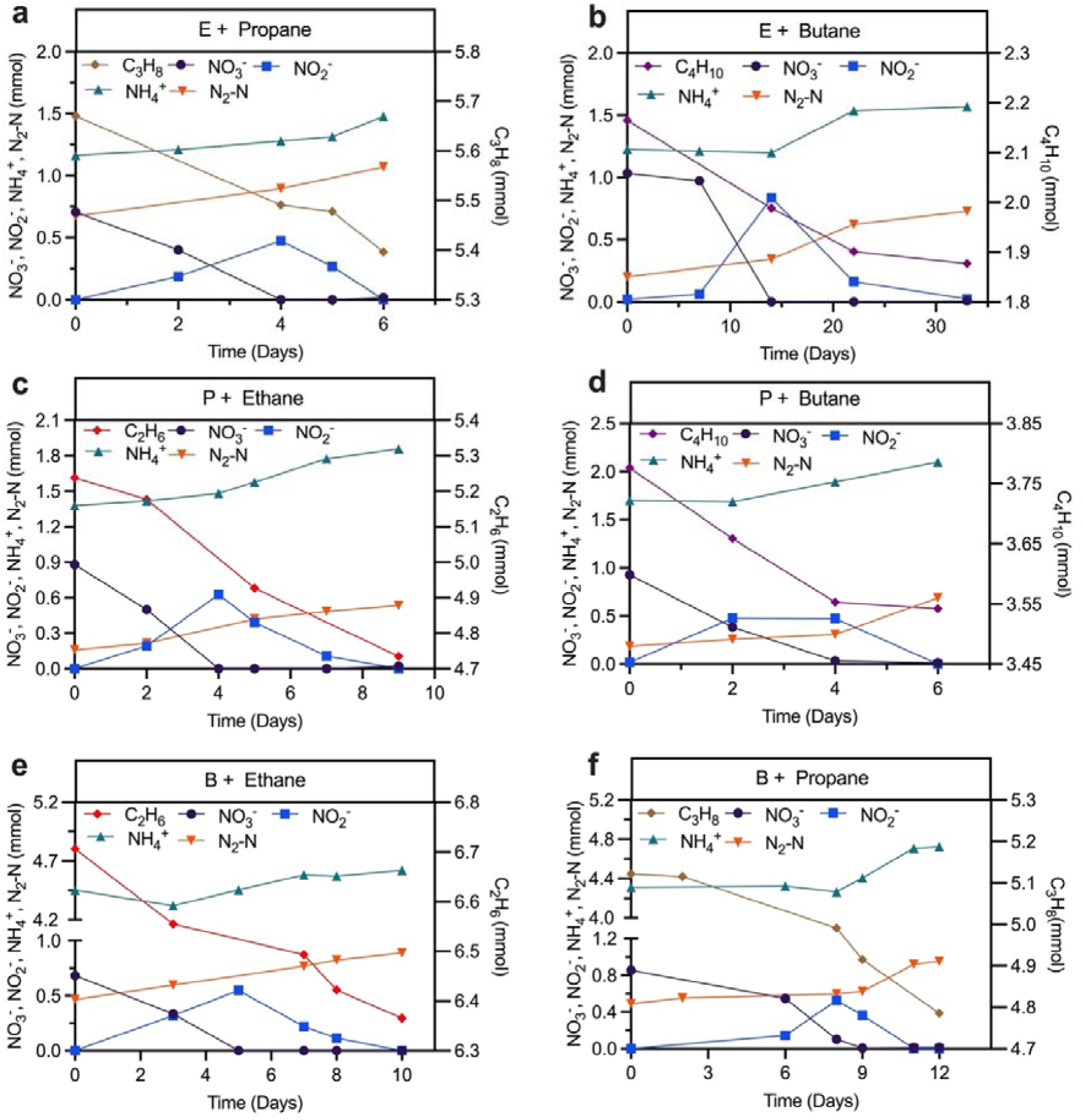
Substrate range tests for the ‘*Ca.* A. nitratireducens’ enriched in the C_2_H_6_-, C_3_H_8_- and C_4_H_10_-fed bioreactors. Subculture from the ethane bioreactor supplemented with propane (**a**) or butane (**b**) showed simultaneous nitrate and propane/butane consumption with production of dinitrogen gas and ammonium. The same was observed for subcultures from the propane bioreactor supplemented with ethane (**c**) and butane (**d**), and the butane bioreactor provided with ethane (**e**) and propane (**f**). Each test was conducted in triplicate (results of other tests were included in Supplementary Fig. 12).

### Implications

This study has identified ‘*Ca*. A. nitratireducens’ as a metabolically diverse anaerobic SCGA oxidiser able to utilise ethane, propane and butane. In previous studies, SRB affiliated with the *Desulfosarcina-Desulfococcus* cluster and the archaeon *Candidatus* ‘Syntrophoarchaeum’ were suggested to be only capable of oxidizing propane and butane, but not ethane^12,13,19^. Conversely, the archaeon *Candidatus* ‘Ethanoperedens thermophilum’ could only oxidize ethane^18^. Importantly, this study is the first to identify a bacterium performing anaerobic ethane oxidation, previously known for archaea only. This study also provides the first physiological evidence for the involvement of the fumarate addition pathway in anaerobic ethane oxidation, closing a key knowledge gap in our understanding of anaerobic SCGA oxidation.

Furthermore, the newly discovered nitrate-dependent anaerobic ethane and butane oxidation (n-DAEO/n-DABO) indicate nitrate is an additional electron sink for C_2_H_6_ and C_4_H_10_, potentially contributing to reducing the negative impacts of C_2_H_6_ and C_4_H_10_ on air quality and on climate. C_2_H_6_ and C_4_H_10_ are recognized as indirect greenhouse gases with net global warming potentials of 10 and 7 times, respectively, that of CO_2_ (100-year horizon) ^33^. Moreover, they also contribute to the production of hazardous substances including carbon monoxide and peroxyacetyl nitrate^34^, which are significant air pollutants. This research advances our understanding of the role of microorganisms in constraining SCGA emissions by identifying another microbially-mediated link between the global carbon and nitrogen cycles. Considering the widespread presence of nitrate and rising emissions of non-methane SCGAs caused by oil and natural gas exploitation^7^, ‘*Ca*. A. nitratireducens’ may play an important role in global carbon and nitrogen cycling.

## Methods

### Bioreactor setup and operation

Activated sludge (50 mL) and anaerobic digestion sludge (100 mL) from a full-scale wastewater treatment plant (Luggage Point, Brisbane, Australia) were used as inoculum for the ethane and n-butane (hereafter butane) bioreactor enrichment. The incubations with ethane and butane were set up in a 1.12 L and a 2.3 L bioreactors, respectively. An anoxic mineral medium^36^ of 0.67 L and 1.69 L was initially added to the ethane and butane reactor, leaving a headspace of 0.3 L and 0.46 L, respectively. The ethane/butane reactors were periodically flushed with pure ethane/butane gas (99.99%, Coregas, Australia) to maintain the ethane/butane partial pressure in the headspace between 0.9 and 1.2 atm. A concentrated stock solution (80 g NO_3_^-^-N l^-1^) was manually pulse-fed to the reactors to replenish NO_3_^-^ to 20–30 mg N/L. The bioreactors were continuously mixed using a magnetic stirrer (IKA, Labtek, Australia) at 650 rpm and operated in a thermostatic chamber (35 ± 1 °C). Every 1–4 months, the stirrers were stopped for 24 h to allow biomass to settle, and the supernatant of 0.2–0.8 L was then replaced with fresh medium. The pH was manually adjusted to 6.8–7.5 using a 1M anoxic HCl solution. Liquid samples (0.4–0.6 mL each) were collected periodically (2–5 samples per week) and filtered immediately using a 0.22 µm membrane filter (polyethersulfone filter, Millex, USA) for the analysis of NO_3_ ^-^, NO_2_ ^-^ and NH_4_ ^+^. A gas sample (100 μL) from the headspace was withdrawn regularly (3–5 times per week) using a gas-tight syringe (1710 SLSYR, Hamilton) for the determination of C_2_H_6_ and N_2_.

### Batch tests for nitrogen and electron balances

Stoichiometric tests were carried out *in-situ* for the biomass of the 1.12 L ethane parent reactor on Days 490, 522 and 559, to investigate nitrogen and electron balances. For stoichiometry determination of nitrate reduction coupled to anaerobic butane oxidation, triplicate batch tests were conducted in 650 mL glass vessels with a subsample of 500 mL biomass anaerobically transferred from the 2.3 L butane parent bioreactor. Total amounts of ethane/butane and N_2_ were calculated by considering ethane/butane/N_2_ in both the headspace (monitored) and liquid phase (calculated with Henry’s law). Two negative control groups were set up in 600 mL bottles: (1) control groups containing only enriched cultures and nitrate (ethane/butane was removed by flushing the bottles with pure argon gas for 20 mins); (2) abiotic control groups without enriched cultures (only synthetic medium containing ethane/butane and nitrate was provided).

### Isotope labelling experiment

A 480 mL sub-culture from the ethane/butane bioreactor was transferred to a 600 mL glass vessel. The ethane culture was flushed with pure C_2_H_6_ for 10 mins, and the 5 mL ^13^C-labelled C_2_H_6_ (^13^CH_3_^13^CH_2_,99 atom % ^13^C, Sigma) was injected into the headspace, followed by an introduction of 0.12 mL nitrate stock solution (40 g N L^-1^) which contained ∼1% ^15^N-labelled NO_3_^-^ (98 atom % ^15^N, Sigma). The butane culture was flushed with argon gas (99.99%, Coregas, Australia) for 20 min. Approximately 24 mL ^13^C-labelled butane (^13^CH_3_^13^CH_2_^13^CH_2_^13^CH_3_, 99 atom% ^13^C, Sigma) was injected into the headspace through the septum. Approximately 1 mL nitrate stock solution (10 g N L^-1^) containing ∼1% ^15^N-labelled sodium nitrate (98 atom % ^15^N, Sigma) was added to achieve a concentration of ∼20 mg N L^-1^. Liquid samples were collected (2–5 samples per week) and filtered through 0.22 µm filters for analysing soluble nitrogen species and respective isotopic fractions. Gaseous samples were collected (4–7 samples in total) from the headspace using a gas-tight syringe (model 1710 SL SYR, Hamilton, USA) and injected into helium-flushed vials (Exetainer, UK) for measuring total C_2_H_6_, C_4_H_10_, CO_2_ and N_2_ in gas phases and their isotopic fractions. For the measurement of the dissolved CO_2_, ∼0.5 mL liquid samples were collected and injected into vacuum vials, followed by acidification with HCl stock solution (1M), and settled for at least 0.5 h to achieve gas-liquid equilibrium before CO_2_ quantification.

### Substrate range tests for ‘*Ca*. A. nitratireducens’

To examine whether the C_2_H_6_-fed culture has the capability of oxidizing propane and butane, two batch tests were set-up by mixing 200 mL culture from the C_2_H_6_-fed bioreactor with 280 mL anoxic mineral medium in 600 mL glass vessels. The two batch reactors were then flushed with pure propane and butane gases, respectively, to remove dissolved ethane and provide propane and butane. The nitrate stock solution (10 g N L^-1^) was added to the reactors to achieve an initial concentration of ∼20 mg N L^-^. The batch tests were conducted in triplicate. Liquid and gas samples were collected as described above. Similarly, cultures from the parent C_3_H_8_- or C_4_H_10_-fed bioreactor were also transferred to new batch reactors and then incubated with ethane and butane, or ethane and propane.

### Chemical analysis

Soluble nitrogen species (NO_3_ ^-^, NO_2_ ^-^, and NH_4_ ^+^) were measured with a flow injection analyser (QuickChem8000, Lachat Instrument, USA). The gas components including C_2_H_6_, CO_2_, and N_2_ in the headspace were determined using a gas chromatograph (GC, 7890A, Agilent, USA) equipped with a Shincarbon ST packed column (Restek, USA) and a thermal conductivity detector (TCD). The GC was operated as described previously^15^.

The butane, ^13^C-labelled butane, ^13^C-labelled ethane, ^13^CO_2_, ^29^N_2_, and ^30^N_2_ in gaseous samples were quantified using a GC (7890A, Agilent, USA) coupled to a quadrupole mass spectrometer (MS, 5957C inert MSD, Agilent, USA). The GC was installed with a J&W HP- PLOT Q PT column (Agilent, USA) using He as the carrier gas at a flow rate of 5.58 mL/min. The GC oven was programmed as follows: (1) samples from the ethane bioreactor: 2 min at 45 ^0^C, ramp at 10 °C/min to 60 ^0^C where it was held for 6 min. (2) samples from the butane bioreactor: 45 °C for 2 min, and then heated with a rate of 15 °C/min to 100 °C where it was hold for 7 min. Mass spectra were detected in the electron impact mode at 70 eV. The mass spectrometer was operated in Selected Ion Monitoring (SIM) mode to detect m/z signals at 30 and 32 Da (C_2_H_6_), 58 and 62 Da (C_4_H_10_), 44 and 45 Da (CO_2_), 28, 29 and 30 Da (N_2_) with a dwell time of 100 ms for each signal. Data processing was performed using the Chemstation program (Agilent, Unite States).

The isotopic fractions of ^15^N-labelled nitrogen-oxyanions (NO_3_^-^ +NO_2_^-^) were analysed using a Thermo Delta V isotope ratio mass spectrometer (IRMS, Thermo Fisher Scientific, USA) following conversion to N_2_O via the denitrifier protocol^37^. In order to measure ^15^N-labelled NO_3_^-^, NO_2_^-^ was removed from the liquid samples with 4% (wt/vol) sulfamic acid in 10% HCl as described previously^38^. The fraction of ^15^N in NO_2_^-^ was calculated according to the difference between ^15^N fraction in nitrogen-oxyanions (NO_3_^-^ +NO_2_^-^) and that in NO_3_^-^. To analyse ^15^N-labelled NH_4_^+^, NH_4_^+^ was trapped in GF/D filters (Whatman, UK) with a microdiffusion method^39^ and then combusted before IRMS analysis.

## 16S rRNA gene amplicon sequencing

Every 2-3 months, 10 mL of biomass samples were taken from the enrichment bioreactors and pelleted by centrifugation (8,000 g for 10 min). DNA extraction was performed using the FastDNA SPIN for Soil kit (MP Biomedicals, USA) according to the manufacture’s protocol. The 16S rRNA gene (V6 to V8 regions) amplicon sequencing was done using the universal primer set 926F (5’-AAACTYAAAKGAATTGACGG-3’) and 1392R (5’-ACGGGCGGTGTGTRC-3’) on an Illumina MiSeq platform (Illumina, USA) at the Australian Centre for Ecogenomics (ACE, Brisbane, Australia). QIIME2 was used to process the sequencing results as described previously^36^.

### Metagenomic sequencing

Biomass collected on Day 746 and 1,150 for ethane and butane bioreactors, respectively, were used for short- and long-read metagenomic sequencing. For short-read sequencing, total DNA was extracted using FastDNA SPIN for Soil kit (MP Biomedicals, USA) and quality controlled using Nanodrop spectrophotometer (Thermo Fisher Scientific, Wilmington, DE) and Qubit^TM^ dsDNA HS Assay Kit. Libraries for short-read sequencing were prepared using Illumina Nextera XT DNA library preparation kit and sequenced on NextSeq 500 (Illumina, USA) platform at ACE.

To obtain Nanopore long reads, total DNA was extracted using Qiagen PowerSoil Pro kit (Qiagen, Germany). Quality of extractions was checked using Qubit 1x dsDNA HS Assay Kit on the Qubit Flex Fluorometer (Thermo Fisher Scientific, Wilmington, DE) and the QIAxcel DNA High Resolution Kit on the QIAxcel Advanced system (Qiagen, Germany). Libraries were prepared and sequenced on PromethION (Oxford Nanopore Technologies, USA).

### Recovery and assessment of microbial populations

Pair-end short reads were trimmed using ReadTrim (https://github.com/jlli6t/ReadTrim) with parameter “--remove_dups --minlen 100”. Nanopore sequencing signals were processed using MinKNOW 20.06.18 and base-called using Guppy 4.0.11 (https://community.nanoporetech.com/), resulting in 53.8 million reads with quality > Q7 with N50 of 2.55kb. Adapters were trimmed using Porechop v0.2.4 (https://github.com/rrwick/Porechop).

Assembly and binning was performed using Aviary (https://github.com/rhysnewell/aviary), which internally called a bunch of different tools, including NanoPack^40^, Flye ^41^, Unicycler^42^, Pilon^43^, Minimap2^44^, CONCOCT^45^, VAMB^46^, MetaBAT 1 & 2^47,48^, MaxBin 2.0^49^ and SemiBin^50^ Specifically, hybrid assembly of short and long reads was performed using workflow ‘assemble’. Resulted assemblies were manually checked using Bandage^51^. Genomes of each community were then recovered using workflow ‘recover’. Obtained genomes were optimized and dereplicated using DASTools 1.1.2^52^. Quality of MAGs was checked using CheckM v1.1.3^53^. Taxonomy information of MAGs was determined using GTDB-Tk 2.1.1^54^. Quality-trimmed short-reads were mapped to assemblies using bowtie 2.3.4.3^55^. Coverage of genome information and other details were viewed and manually checked using IGV 2.11.1^56^. Abundance of each MAG was profiled using CoverM 0.6.1 (https://github.com/wwood/CoverM). Genome characteristics were calculated using BioSut (https://github.com/jlli6t/BioSut).

### Functional annotation

Preliminary annotation across MAGs and unbin contigs were performed using Prokka 1.14.5^57^. Predicted protein sequences were then searched against KEGG (July 2021) using kofamscan 1.3.0^58^, the hit with e-value < 1e-10 and maximal F-measure was selected for each gene. UniRef100^59^ (March 2020) was searched against using diamond^60^ v2.0.11.149 with ‘blastp --sensitive’. The best hit with e-value < 1e-5 and identity > 30 was selected for each gene and mapped to the KEGG Orthology database. The eggNOG v5^61^ was searched against using emapper 2.1.5^62^. Metabolic pathways were reconstructed using KEGG. Pathways identified to be > 75% complete were considered as ‘expressed’. Full-length AssA genes from the P and B MAGs were identified based on blastn hits to the AssA genes from E MAG and translated using NCBI’s ORF finder.

### Phylogenetic analysis of recovered Symbiobacteriia genomes

*Genome tree.* Phylogenetic placement of the two recovered Symbiobacteriia genomes in current study was performed with the existing ‘*Ca.* A. nitratireducens’ MAG^15^ and available Firmicutes genomes in GTDB r207^63,64^ using 120 bacterial-specific conserved marker genes. Briefly, marker genes in genomes were identified using Prodigal 2.6^65^ and aligned using HMMER 3.3^66^. Trees were inferred using FastTree 2.1.11^67^ with WAG+GMMA models. Bootstrap of the constructed tree was performed using workflow ‘bootstrap’ from GenomeTreeTk v0.1.6 (https://github.com/dparks1134/GenomeTreeTk) with 100 times nonparametric bootstrapping. The tree was visualized using ARB 6.0.6^68^ and further refined using Adobe Illustrator (Adobe, USA).

*AssA amino acid tree.* The different AssAs in ‘*Ca.* A. nitroreducens’ were aligned with reference AssA, BssA and MasD protein sequences downloaded from Uniprot database using muscle 3.8.31^69^. We applied trimAI 1.4.1^70^ to trim gaps in msa. FastTree 2.1.11^67^ was used to infer the phylogenetic tree. Bootstrap value was calculated, and tree was visualized as per the genome tree construction.

### Metatranscriptomic sequencing and data analysis

Two distinct phases were observed for the nitrate reduction in both ethane and butane bioreactors (Fig. 1a, 1b). For the total RNA extraction, the active enriched culture (10 mL) collected from each phase was mixed with 30 mL of RNAlater solution (Sigma-Aldrich) and left to stand for 1h before extraction. Total RNA was then extracted using the RNeasy Powersoil Total RNA kit (Qiagen, Germany) according to the manufacturer’s protocols. Removal of genomic DNA contamination was performed using a Turbo DNA-free kit (Thermo Fisher Scientific, USA), followed by concentration with a RNA Clean & Concentrator-5 Kit (Zymo Research, USA). RNA library was prepared using the TruSeq Total RNA Library Prep with Ribo-Zero Plus kit following the manufacture’s protocol. The library was sequenced on a NovSeq6000 (Illumina, USA) platform at ACE (Brisbane, Australia) in 2 × 75 cycles paired-end runs.

The metatranscriptomic paired-end reads were mapped to dereplicated genome sets and filtered using minimum cut-off values of 97% identity and 75% alignment. The Symbiobacteriia MAG generated from the ethane system was selected as the representative SymBio MAG due to the presence of full length AssA genes. TranscriptM (GitHub - sternp/transcriptm) was used to unambiguously mapped mRNA for each ORF and calculate the total transcripts per million (TPM).

### Protein extraction and metaproteomics

For protein extraction, enrichment cultures collected from Phase 1 and 2 (10 mL each phase) were pelleted by centrifugation (18,000 g, 4 °C) and then washed with 1 × PBS. The cell lysis was performed by adding 5% sodium dodecyl sulfate (SDS) and then incubating with 20 mM dithiothreitol (final concentration) at 70 °C for 1 h. After cooling to room temperature, the protein solution was alkylated with 40 mM iodoacetamide (final concentration) in the dark for 0.5 h. Afterwards, 1.2% phosphoric acid (final concentration) and six volumes of S-Trap binding buffer (90% methanol, 100 mM final concentration of ammonium bicarbonate, pH 7.1) were added. Total protein was digested in a S-Trap Micro Spin Column (ProtiFi, Huntington, USA) as described previously^15^. The digested peptides were analysed by Liquid Chromatography-tandem Mass Spectrometry (LC-MS/MS) using a Dionex Ultimate 3000 RSLCnano-LC system coupled to a Q-Exactive^TM^ H-X Hybrid Quadrupole-Orbitrap™ mass spectrometer (Thermo Scientific^TM^). Raw sequencing data were searched against the annotated closed genomes of Strain E and B, respectively, in Thermo Proteome Discoverer. The identified proteins contained at least 1 unique peptide with a stringency cut-off of false discovery rate (FDR, *q* value) less than 0.05.

### Metabolite extraction and detection

For metabolite extractions, enrichment cultures (5 mL) collected from ethane and butane bioreactors were centrifuged at 10,000 rpm for 10 min (4 °C) to harvest the cells. The metabolites were extracted from pelleted cells as described previously^15^. The ethyl and butyl succinate standards (custom synthesized by Best of Chemicals, USA) and cell extracts were then processed and analysed using an ultra-high-sensitivity triple quadrupole GC/MS-50 system (Shimadzu, Japan). The three most abundant fragmentation ions were chosen to monitor, with Transient 1 used as quantifier and the other two as qualifiers (See Supplementary Table 9).

### Computational analyses for catalytic subunits of different alkylsuccinate synthases (AssAs) in ‘*Ca*. A. nitratireducens’

#### Structural Modelling and Molecular Dynamics (MD) Simulation

The amino acid sequences of three complete AssAs were acquired from the closed genome of ‘*Ca*. A. nitratireducens’ in the ethane-fed system. The tertiary structures of AssAs were modelled with Alphafold-2^71^. Fumarate and alkanes were bound to corresponding AssA by CB-dock-2^72^. AssA-Fumarate-Alkane complexes were solvated by CHARMM-GUI^73^ with a thickness of 15 Å. Water type was TIP3P^74^ and the force field was CHARMM36m^75^. NaCl (200 mM) was used to ionize the systems^76^. The final systems were then subjected to the MD simulations with NAMD 2.12^77^. Periodic boundary condition was applied to the simulating box, and particle mesh Ewald was used for the long-range electrostatic interactions. The pressure was set at 1 atm using a Langevin thermostat with a damping coefficient of 1/ps. A Nose−Hoover Langevin piston barostat with a decay period of 25 fs was applied. The temperature was reassigned every 500 steps. Simulations for each model include two steps. The first is 1 ns equilibration (NVP), and the second is 50 ns production run (NPT).

#### Root mean square fluctuation (RMSF) and deviation (RMSD) calculations

The RMSF for α-carbons of the amino acid residues is calculated with equation 1^78,79^.

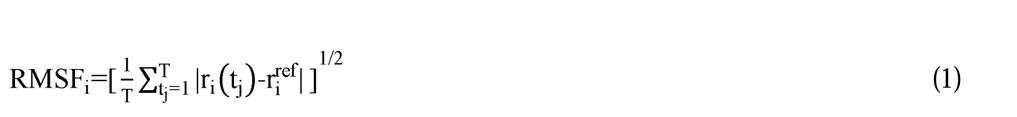

where i represents the residue ID, T represents the total simulation time (Here is the number of frames) and r_i_(t_j_) represents the position of residues i at time t_j_. The r ^ref^ is the reference position of residue i, calculated by the time-average position.

To measure the average distance between two protein structures, the RMSD is calculated with equation 2

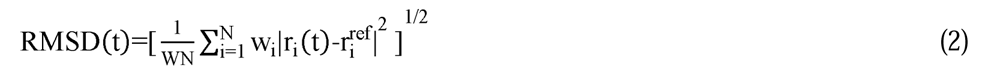

where W = Σw_i_ is the weighting factor, and N is the total number of atoms. The r_i_(t) is the position of atom i at time t after least square fitting the structure to the reference structure. The r_i_^ref^ is the reference position of residue i defined by the reference structure (Here we used the initial structure as the reference).

#### Hydrogen bond analyses

The hydrogen bonds were analysed by VMD^79^ based on 100 frames obtained from the last MD simulations of 40 to 50 ns. The cut-off distance and angle for hydrogen bond analyses were set as 3.5 Å and 20°, respectively.

## Data availability

Sequencing data are archived in NCBI database under Project number PRJNA989758. The mass spectrometry proteomics data have been deposited to the ProteomeXchange Consortium via the PRIDE partner repository with the dataset identifier PXD039267.

## Supporting information

SI data 1-MAG information

## Acknowledgements

We acknowledge G. Talbo for help with protein extraction and proteomics data analysis, T. Stark for assistance with isotope carbon and metabolite analyses, N. Dawson, J. Li for chemical analyses and J.P. Engelberts for assistance with FISH. J.L. is supported by UQ Research Training Scholarship. J.G., S.M. and G.T. are supported by Australian Research Council (ARC) Future Fellowships FT170100196, FT190100211 and FT170100070, respectively. Z.Y. is supported by ARC Australian Laureate Fellowship (FL170100086).

## Author Contributions

J.G. conceived the study. M.W., J.L., C.L., J.G. and S.M. planned the experiments. M.W. run the butane-fed bioreactor and performed mass and electron balance and isotope labelling experiments. C.L. run the ethane-fed bioreactor and performed mass and electron balance and isotope labelling experiments. J.L., A.L. and M.W. performed the bioinformatics analysis, with support from S.M., G.T., C.L. and J.G. D.E. conducted the isotope nitrogen measurements. S.M. completed FISH microscopy. M.W. and C.L. performed the sampling, preservation, DNA, RNA and protein extractions for metagenomics, metatranscriptomics and metaproteomics sequencing. L.L. set up methods for the protein extractions. S.S., M.W. and L.L. performed the structural modelling and molecular dynamics simulations. M.W. and R.G. conducted the substrate range test experiments. M.W., C.L., J.G. and Z.Y. performed the process data analysis. M.W., J.L, C.L. and J.G. wrote the manuscript in consultation with all other authors.

