## Supporting information for "Nitrate-driven anaerobic oxidation of ethane and butane by bacteria"

**1. Supplementary Tables**

**Supplementary Table 1.** Nitrogen and electron balances for the enrichment cultures capable of coupling anaerobic ethane/butane oxidation to nitrate reduction in the batch tests.

| Nitrogen and electron balance (mmol/L) | C_2_H_6_/  C_4_H_10_ oxidized | NO_3_^-^ and NO_2_^-^reduced | NH_4_^+^ generated | N_2_-N generated | Maximum electrons available from C_2_H_6_/C_4_H_10_ oxidation | Electrons required for NO_3_^-^reduction^*^ | Nitrogen balance^†^ | Electron balance^‡^ |
| --- | --- | --- | --- | --- | --- | --- | --- | --- |
| Ethane bioreactor | 0.89 | 1.78 | 0.58 | 1.37 | 12.44 | 11.43 | 0.92 | 1.09 |
|  | 1.02 | 1.86 | 1.08 | 1.10 | 14.26 | 14.14 | 0.86 | 1.01 |
|  | 0.91 | 1.68 | 0.79 | 0.88 | 12.72 | 10.74 | 1.00 | 1.18 |
| Butane bioreactor | 0.49 | 1.74 | 0.96 | 0.73 | 12.77 | 11.31 | 1.03 | 1.13 |
|  | 0.34 | 1.43 | 0.71 | 0.77 | 8.84 | 9.56 | 0.97 | 0.92 |
|  | 0.36 | 1.71 | 0.74 | 0.69 | 9.34 | 9.41 | 1.19 | 0.99 |

^*^ Electrons required for NO_3_^-^  reduction = NH_4_^+^ generated $\times$8 + N_2_-N generated $\times$5.

^†^ Nitrogen balance = (NO_3_^-^  reduced)/ (NH_4_^+^ generated + N_2_-N generated ).

^‡^ Maximum electrons available from ethane (=C_2_H_6_ oxidized $\times$14) or butane (=C_4_H_10_ oxidized $\times$26) oxidation divided by electrons required for NH_4_^+^ and N_2_-N production; theoretically higher than 1.0, due to a fraction of carbon assimilated into biomass cells.

**Supplementary Table 2. Sequencing statistics of metagenomics and metatranscriptomics.** Key features of the metagenome (Illumina short-read and Nanopore long-read sequencing) and metatranscriptomic datasets generated for the C_2_H_6_- and C_4_H_10_-fed enrichment cultures. Two samples were taken from the C_2_H_6_-fed bioreactors for short-read sequencing, thus generating two datasets.

|  | |  | | **C_2_H_6_-fed cultures** | | | | **C_4_H_10_-fed cultures** |
| --- | --- | --- | --- | --- | --- | --- | --- | --- |
| **Metagenomic short read sequencing** | | # trimmed reads | | 99,139,736/102,953,198 | | | | 116,463,546 |
|  |  | % Reads in metagenomic scaffolds | | 94.1/94.5 | | | | 96.5 |
| **Long-read sequencing** | | # raw nanopore reads (million) | | 11.8 | | | 28.9 | |
|  |  | Maximum read length | | 310,518 | | | 658,543 | |
|  |  | N50 | | 4,440 | | | 544 | |
| **Hybrid assembly** | | Assembled metagenome size (Mbp) | | 474.6 | | | | 314.6 |
|  |  | N50 (bp) | | 22,096 | | | | 100,544 |
|  |  | Maximum scaffold (bp) | | 6,053,062 | | | | 4,746,968 |
|  |  | # scaffold | | 100,732 | | | | 22,756 |
| **Metatranscriptome** | **Total Illumina reads** | | **Trimmed Reads** | | **non_rRNA reads** | **mRNA mapped to MAGs** | | |
| C2-phase1-RNA | 125,039,398 | | 116,358,434 | | 82,846,667 | 34,950,915 | | |
| C2-phase2-RNA | 212,602,712 | | 194,685,456 | | 137,965,946 | 71,243,694 | | |
| C4-phase1-RNA | 68474852 | | 64,488,864 | | 40,744,382 | 25,927,776 | | |
| C4-phase2-RNA | 62856468 | | 59,114,204 | | 37,435,066 | 23,625,080 | | |

**Supplementary Table 3. Estimated abundance of each lineage recovered from C_2_H_6_-fed community.** Lineages with abundance over 1% were shown in the table.

| **Bin ID** | **C2_ Phase1-DNA (%)** | **C2_ Phase2-DNA (%)** | **Classification** |
| --- | --- | --- | --- |
| C2_01 | 28.46 | 26.34 | d__Bacteria;p__Patescibacteria;c__4484-211;o__4484-211;f__;g__;s__ |
| **C2_SYM** | **14.38** | **15.55** | **d__Bacteria;p__Firmicutes_E;c__Symbiobacteriia;o__;f__;g__;s__** |
| C2_02 | 10.75 | 11.29 | d__Bacteria;p__Armatimonadota;c__Fimbriimonadia;o__Fimbriimonadales;f__Fimbriimonadaceae;g__OLB18;s__OLB18 sp001567425 |
| C2_03 | 6.77 | 7.56 | d__Bacteria;p__Bacteroidota;c__Ignavibacteria;o__Ignavibacteriales;f__Melioribacteraceae;g__DSXH01;s__ |
| C2_04 | 3.23 | 3.63 | d__Bacteria;p__Chloroflexota;c__Anaerolineae;o__Promineofilales;f__Promineofilaceae;g__Promineofilum;s__ |
| C2_05 | 3.83 | 3.30 | d__Bacteria;p__Bacteroidota;c__Bacteroidia;o__Flavobacteriales;f__Vicingaceae;g__BCD5;s__BCD5 sp013112825 |
| C2_06 | 2.36 | 2.74 | d__Bacteria;p__Proteobacteria;c__Gammaproteobacteria;o__Burkholderiales;f__SG8-39;g__;s__ |
| C2_07 | 2.46 | 2.45 | d__Bacteria;p__Actinobacteriota;c__Acidimicrobiia;o__ATN3;f__ATN3;g__ATN3;s__ |
| C2_08 | 2.12 | 1.94 | d__Bacteria;p__Chloroflexota;c__Anaerolineae;o__Anaerolineales;f__EnvOPS12;g__UBA7227;s__UBA7227 sp002473085 |
| C2_09 | 1.91 | 1.67 | d__Bacteria;p__Patescibacteria;c__Microgenomatia;o__UBA1406;f__GWC2-37-13;g__GWC2-37-13;s__GWC2-37-13 sp002050095 |
| C2_10 | 1.51 | 1.48 | d__Bacteria;p__Bacteroidota;c__Ignavibacteria;o__Ignavibacteriales;f__Ignavibacteriaceae;g__IGN2;s__IGN2 sp013285405 |
| C2_11 | 1.19 | 1.37 | d__Bacteria;p__Chloroflexota;c__Anaerolineae;o__Anaerolineales;f__EnvOPS12;g__OLB14;s__ |
| C2_12 | 1.06 | 1.28 | d__Bacteria;p__Acidobacteriota;c__Vicinamibacteria;o__Vicinamibacterales;f__Fen-181;g__;s__ |
| C2_13 | 1.26 | 1.10 | d__Bacteria;p__Bacteroidota;c__UBA10030;o__UBA10030;f__;g__;s__ |

**Supplementary Table 4. Estimated abundance of each lineage recovered from C_4_H_10_-fed community.** Lineages with abundance over 1% were shown in the table.

| **Bin ID** | **C4_DNA (%)** | **Classification** |
| --- | --- | --- |
| **C4_SYM** | **16.72** | **d__Bacteria;p__Firmicutes_E;c__Symbiobacteriia;o__;f__;g__;s__** |
| C4_01 | 15.46 | d__Bacteria;p__Chloroflexota;c__Anaerolineae;o__Promineofilales;f__Promineofilaceae;g__JAAYYY01;s__JAAYYY01 sp012515125 |
| C4_02 | 15.39 | d__Bacteria;p__Planctomycetota;c__Phycisphaerae;o__Phycisphaerales;f__SM1A02;g__CAADGN01;s__CAADGN01 sp900696545 |
| C4_03 | 11.67 | d__Bacteria;p__Chloroflexota;c__Anaerolineae;o__Anaerolineales;f__Anaerolineaceae;g__Bellilinea;s__ |
| C4_04 | 10.50 | d__Bacteria;p__Chloroflexota;c__Anaerolineae;o__Anaerolineales;f__EnvOPS12;g__UBA7227;s__UBA7227 sp002473085 |
| C4_05 | 6.63 | d__Bacteria;p__Proteobacteria;c__Alphaproteobacteria;o__Rhizobiales;f__Rhizobiaceae;g__Aquamicrobium_A;s__ |
| C4_06 | 3.95 | d__Bacteria;p__Chloroflexota;c__Anaerolineae;o__Anaerolineales;f__Anaerolineaceae;g__Bellilinea;s__ |
| C4_07 | 1.33 | d__Bacteria;p__Bacteroidota;c__Ignavibacteria;o__Ignavibacteriales;f__Ignavibacteriaceae;g__Ignavibacterium;s__Ignavibacterium sp900696555 |
| C4_08 | 1.25 | d__Bacteria;p__Proteobacteria;c__Gammaproteobacteria;o__Burkholderiales;f__Burkholderiaceae;g__QOAZ01;s__ |
| C4_09 | 1.23 | d__Bacteria;p__Patescibacteria;c__Microgenomatia;o__UBA1406;f__GWC2-37-13;g__GWC2-37-13;s__GWC2-37-13 sp002050095 |
| C4_10 | 1.16 | d__Bacteria;p__Proteobacteria;c__Alphaproteobacteria;o__Rhizobiales;f__Xanthobacteraceae;g__Palsa-892;s__ |
| C4_11 | 1.14 | d__Bacteria;p__Proteobacteria;c__Gammaproteobacteria;o__Burkholderiales;f__Burkholderiaceae;g__SCN-69-89;s__SCN-69-89 sp001724855 |

**Supplementary Table 5. Calculated expression level of each genome in C_2_H_6_-fed bioreactor.** Genomes with expression level >1% in at least one phase were shown**.**

| **Bin ID** | **C2_Phase1-RNA (%)** | **C2_Phase2-RNA (%)** |
| --- | --- | --- |
| C2_SYM | 62.58 | 60.35 |
| C2_02 | 13.01 | 9.92 |
| C2_04 | 3.47 | 4.29 |
| C2_06 | 2.51 | 2.58 |
| C2_11 | 2.68 | 3.81 |
| C2_12 | 2.66 | 2.94 |
| C2_17 | 0.84 | 1.04 |
| C2_26 | 1.60 | 1.77 |

**Supplementary Table 6. Calculated expression level of each genome in C_4_H_10_-fed bioreactor.** Genomes with expression level >1% in at least one phase were shown**.**

| **Bin ID** | **C4_Phase1-RNA (%)** | **C4_Phase2-RNA (%)** |
| --- | --- | --- |
| C4_SYM | 84.70 | 84.27 |
| C4_01 | 2.89 | 3.59 |
| C4_02 | 5.09 | 3.32 |
| C4_03 | 1.34 | 1.23 |
| C4_04 | 2.45 | 3.54 |

**Supplementary Table 7.** Average amino acid identities (AAI) between different AssAs in ‘*Ca.* A. nitratireducens’ genomes recovered from the ethane- (E), propane- (P) and butane-fed (B) bioreactors. AAI values were calculated using Blastp.

| **AAI** | **B_AssA1** | **B_AssA2** | **B_AssA3** | **E_AssA1** | **E_AssA2** | **E_AssA3** | **P_AssA1** | **P_AssA2** | **P_AssA3** |
| --- | --- | --- | --- | --- | --- | --- | --- | --- | --- |
| **B_AssA1** | - | 89.66 | 61.95 | 100.00 | 90.96 | 90.73 | 100.00 | 93.10 | 90.74 |
| **B_AssA2** | 89.66 | - | 91.43 | 89.66 | 95.93 | 97.81 | 89.66 | 95.94 | 97.81 |
| **B_AssA3** | 61.95 | 91.43 | - | 61.95 | 96.91 | 100.00 | 61.95 | 96.24 | 100.00 |
| **E_AssA1** | 100.00 | 89.66 | 61.95 | - | 90.96 | 90.73 | 100.00 | 93.10 | 90.73 |
| **E_AssA2** | 90.96 | 95.93 | 96.91 | 90.96 | - | 96.60 | 90.96 | 100.00 | 96.60 |
| **E_AssA3** | 90.73 | 97.81 | 100.00 | 90.73 | 96.60 | - | 90.73 | 96.24 | 100.00 |
| **P_AssA1** | 100.00 | 89.66 | 61.95 | 100.00 | 90.96 | 90.73 | - | 93.10 | 90.74 |
| **P_AssA2** | 93.10 | 95.94 | 96.24 | 93.10 | 100.00 | 96.24 | 93.10 | - | 96.24 |
| **P_AssA3** | 90.74 | 97.81 | 100.00 | 90.73 | 96.60 | 100.00 | 90.74 | 96.24 | - |

**Supplementary Table 8.** Average nucleotide identities (ANI) between different AssAs in ‘*Ca.* A. nitratireducens’ genomes recovered from the ethane- (E), propane- (P) and butane-fed (B) bioreactors. ANI values were calculated using Blastn.

| **AAI** | **B_AssA1** | **B_AssA2** | **B_AssA3** | **E_AssA1** | **E_AssA2** | **E_AssA3** | **P_AssA1** | **P_AssA2** | **P_AssA3** |
| --- | --- | --- | --- | --- | --- | --- | --- | --- | --- |
| **B_AssA1** | - | 89.31 | 89.57 | 99.96 | 89.85 | 89.57 | 99.96 | 89.78 | 89.57 |
| **B_AssA2** | 89.31 | - | 98.83 | 89.27 | 97.42 | 98.87 | 89.27 | 97.46 | 98.87 |
| **B_AssA3** | 89.57 | 98.83 | - | 89.61 | 96.80 | 99.96 | 89.61 | 96.65 | 99.96 |
| **E_AssA1** | 99.96 | 89.27 | 89.61 | - | 89.89 | 89.61 | 100.00 | 89.74 | 89.61 |
| **E_AssA2** | 89.85 | 97.43 | 96.80 | 89.89 | - | 96.84 | 89.89 | 99.84 | 96.84 |
| **E_AssA3** | 89.57 | 98.87 | 99.96 | 89.61 | 96.84 | - | 89.61 | 96.68 | 100.00 |
| **P_AssA1** | 99.96 | 89.27 | 89.61 | 100.00 | 89.89 | 89.61 | - | 89.74 | 89.61 |
| **P_AssA2** | 89.78 | 97.47 | 96.65 | 89.74 | 99.84 | 96.69 | 89.74 | - | 96.69 |
| **P_AssA3** | 89.57 | 98.87 | 99.96 | 89.61 | 96.84 | 100.00 | 89.61 | 96.68 | - |

**Supplementary Table 9.** Optimized multiple reaction monitoring transitions and retention times for ethylsuccinate and butylsuccinate analysed by GC-MS/MS.

| Compound | Retention time (min) | Transient 1 (m/z) | Collision Energy (V) | Transient 2 (m/z) | Collision Energy (V) | Transient 3 (m/z) | Collision Energy (V) |
| --- | --- | --- | --- | --- | --- | --- | --- |
| Ethyl succinate | 9.940 | 275.0>73.1 | 35 | 217.0>55.1 | 10 | 275.0>147.2 | 10 |
| Butyl succinate | 12.245 | 303.0>147.1 | 10 | 147.1>73.1 | 30 | 231.0>69 | 10 |

**2. Supplementary Figures**

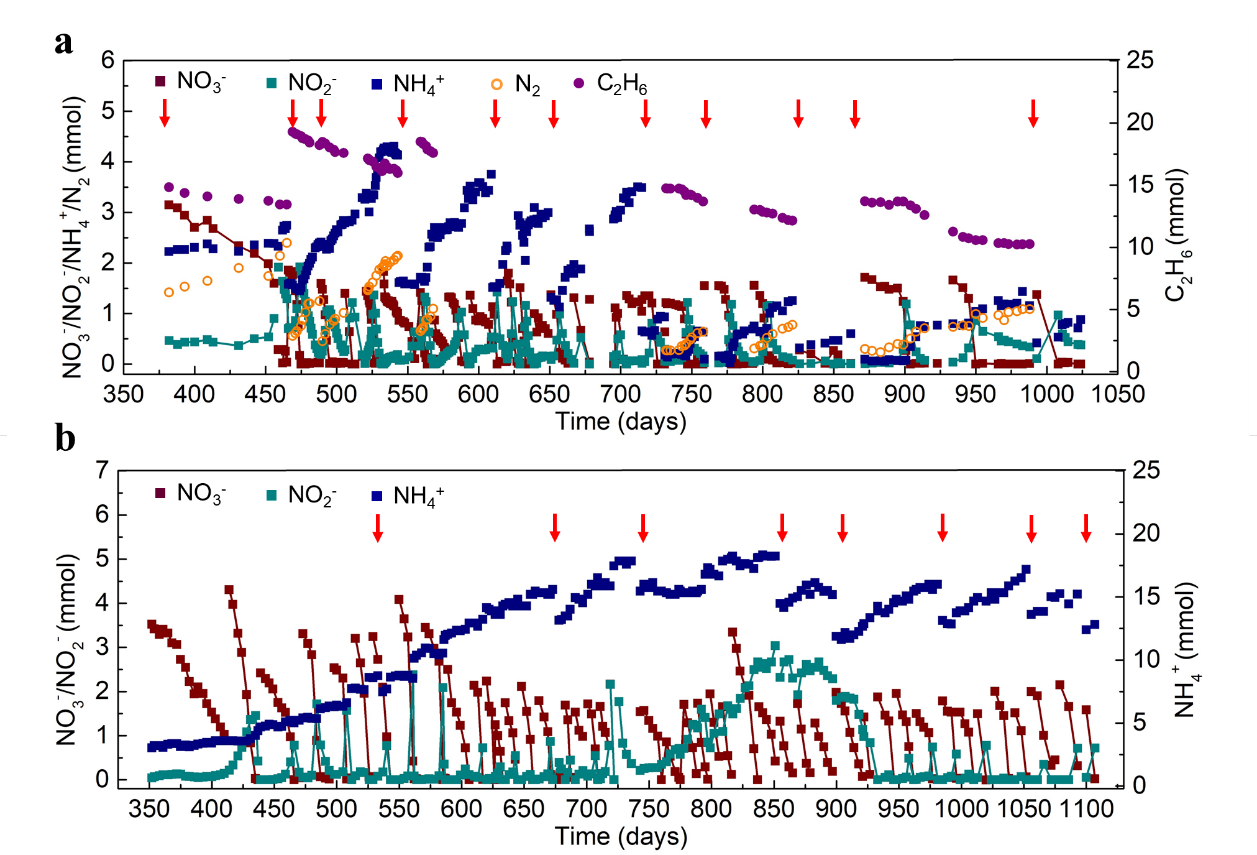

**Supplementary Fig. 1** Long-term performance of the C_2_H_6_- **(a)** and C_4_H_10_-fed **(b)** bioreactors. **(a)** Simultaneous C_2_H_6_ and NO_3_^-^ consumption with transitory formation of NO_2_^-^, and production of N_2_ and NH_4_^+^ in the C_2_H_6_-fed bioreactor. **(b)** NO_3_^-^ consumption with production of NH_4_^+^ and transitory formation of NO_2_^-^ was also observed in the C_4_H_10_-fed bioreactor. Red arrows indicate medium replacements and the flushing of the bioreactor headspace with C_2_H_6_ or C_4_H_10_.

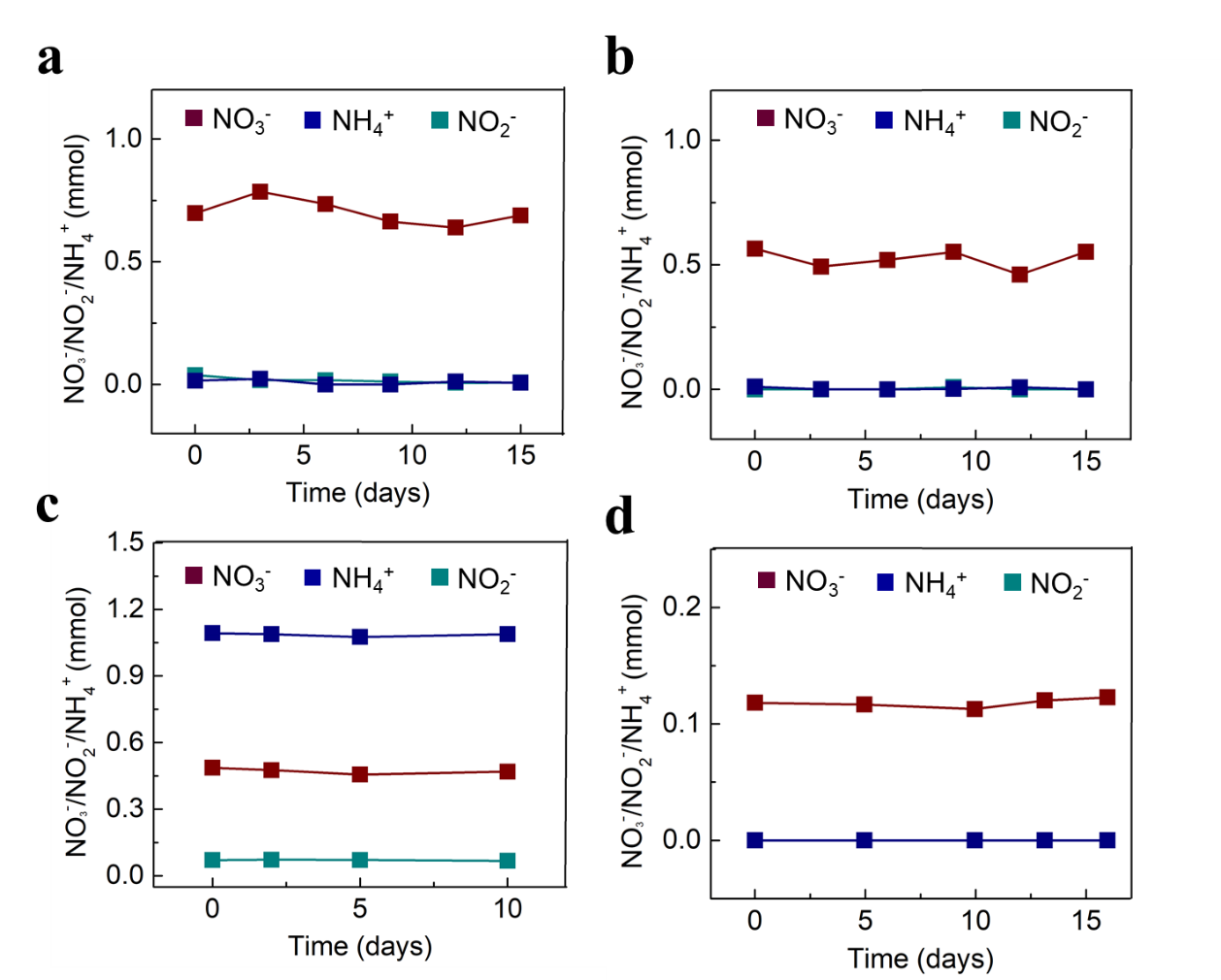

**Supplementary Fig. 2 Profiles of NO_3_^-^, NO_2_^-^ and NH_4_^+^ in the control tests.** **a, c**, negligible NO_3_^-^ consumption or NO_2_^-^/NH_4_^+^ production in the absence of C_2_H_6_ **(a)** or C_4_H_10_ **(c)**. **b, d,** negligible NO_3_^-^ consumption or NO_2_^-^/NH_4_^+^ production in the abiotic control (without biomass) supplied with C_2_H_6_ **(b)** or C_4_H_10_ **(d)**.

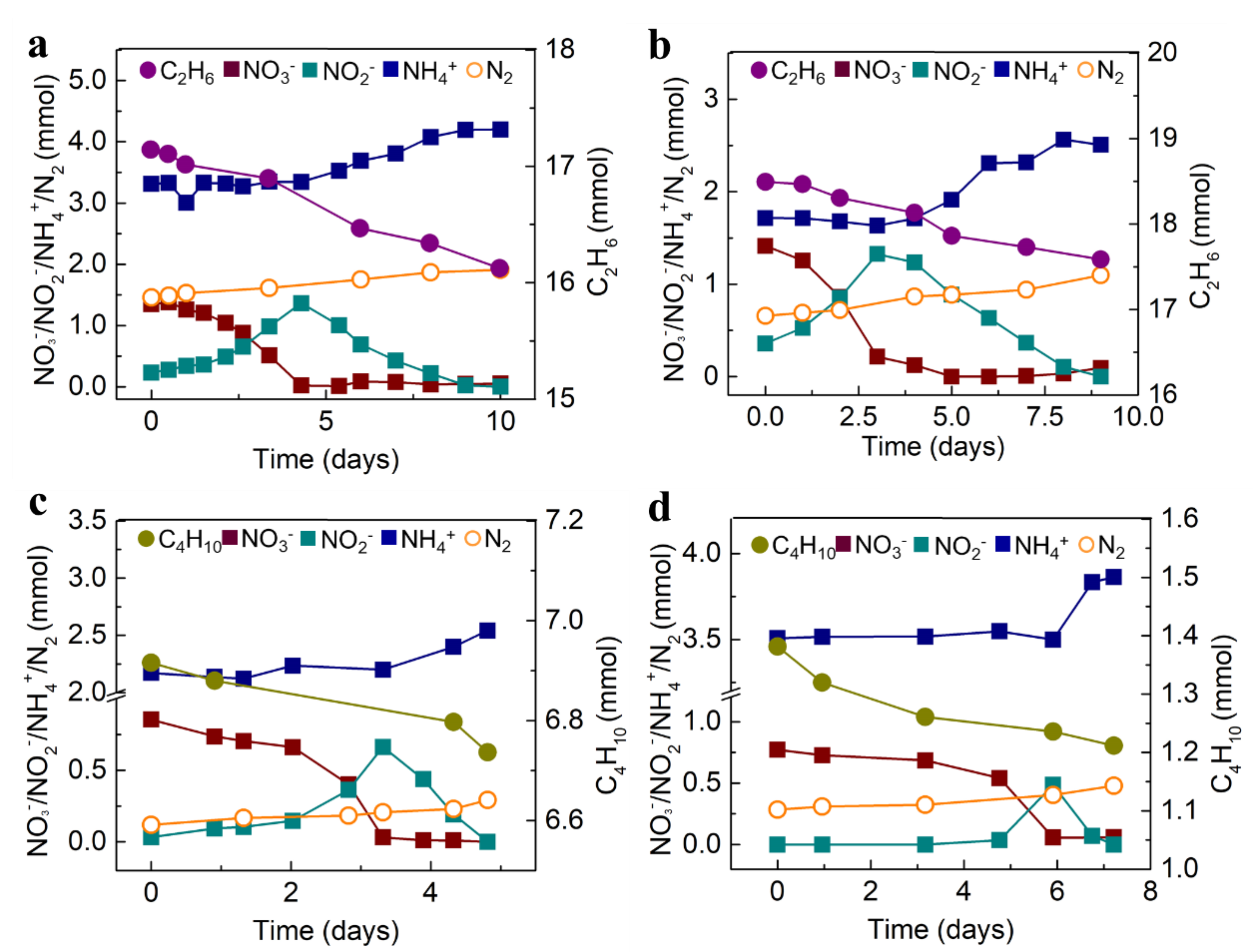

**Supplementary Fig. 3 Profiles of C_2_H_6_, C_4_H_10_, and nitrogen species in the batch tests. a, b,** supplementary batch tests for C_2_H_6_-fed bioreactor (started on Day 522 and 559). **c, d,** supplementary batch tests for C_4_H_10_-fed bioreactor (started on Day 1,100 and 1,124).

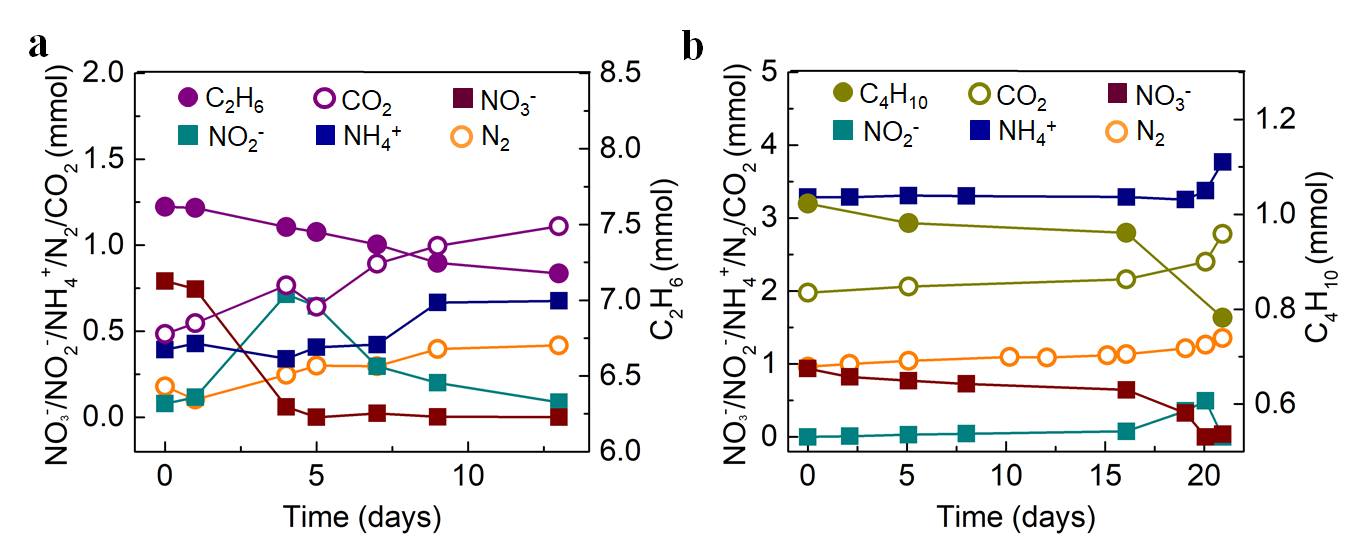

**Supplementary Fig. 4 Profiles of total C_2_H_6_, C_4_H_10_, and nitrogen species during isotope labelling tests. a, b,** oxidation of C_2_H_6_ (**a**) or C_4_H_10_ **(b**) to CO_2_, and reduction of NO_3_^-^ to NH_4_^+^ and N_2_ with temporary accumulation of NO_2_^-^.

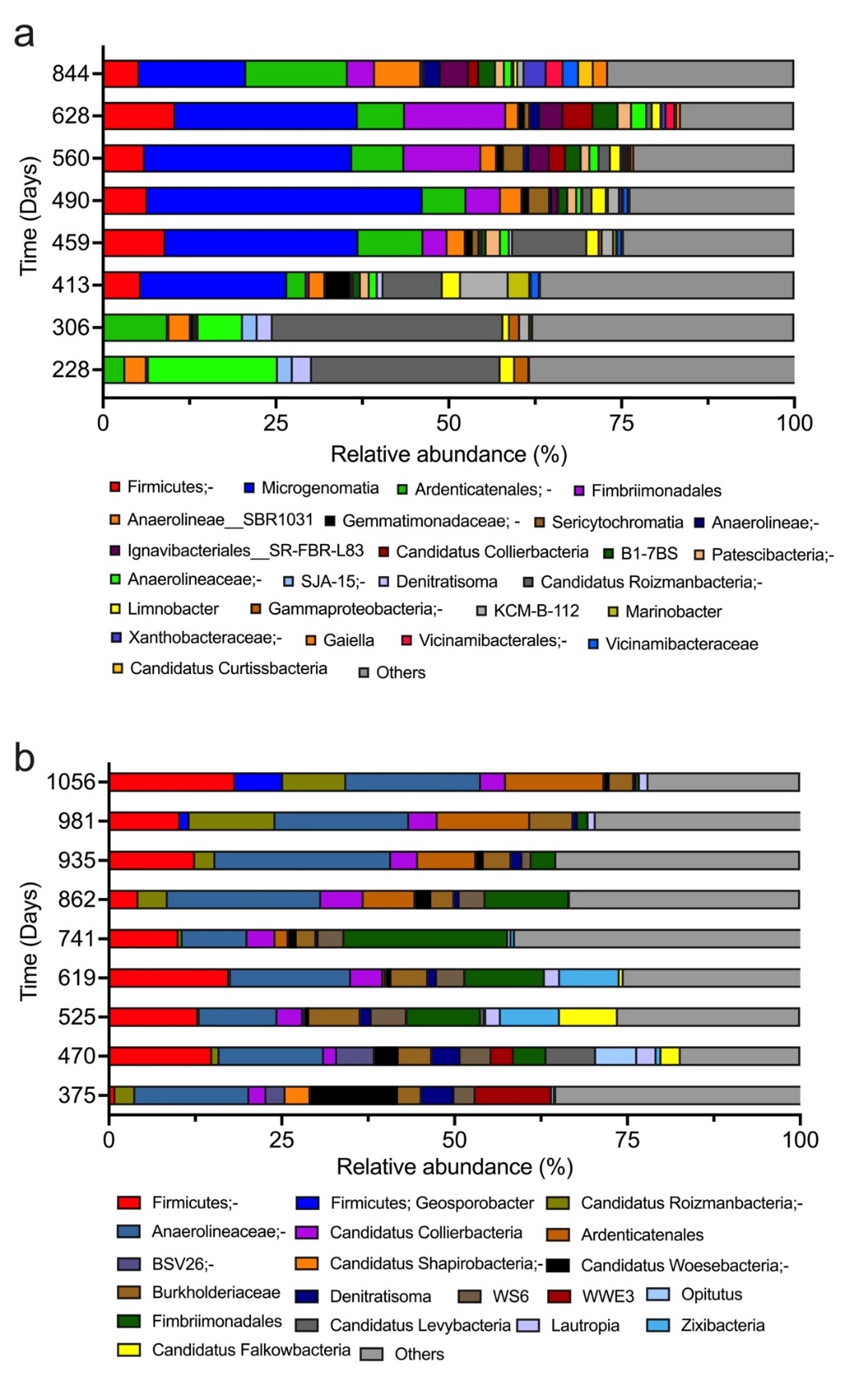

**Supplementary Fig. 5** Phylotypes in the enrichment cultures at the genus level based on 16S rRNA gene amplicon sequencing during long-term operations of C_2_H_6_- **(a)** and C_4_H_8_-fed **(b)** bioreactors. Genera with an abundance of ≥2% in at least one sample are displayed, while genera account for <2% in all samples are classified as ‘Others’.

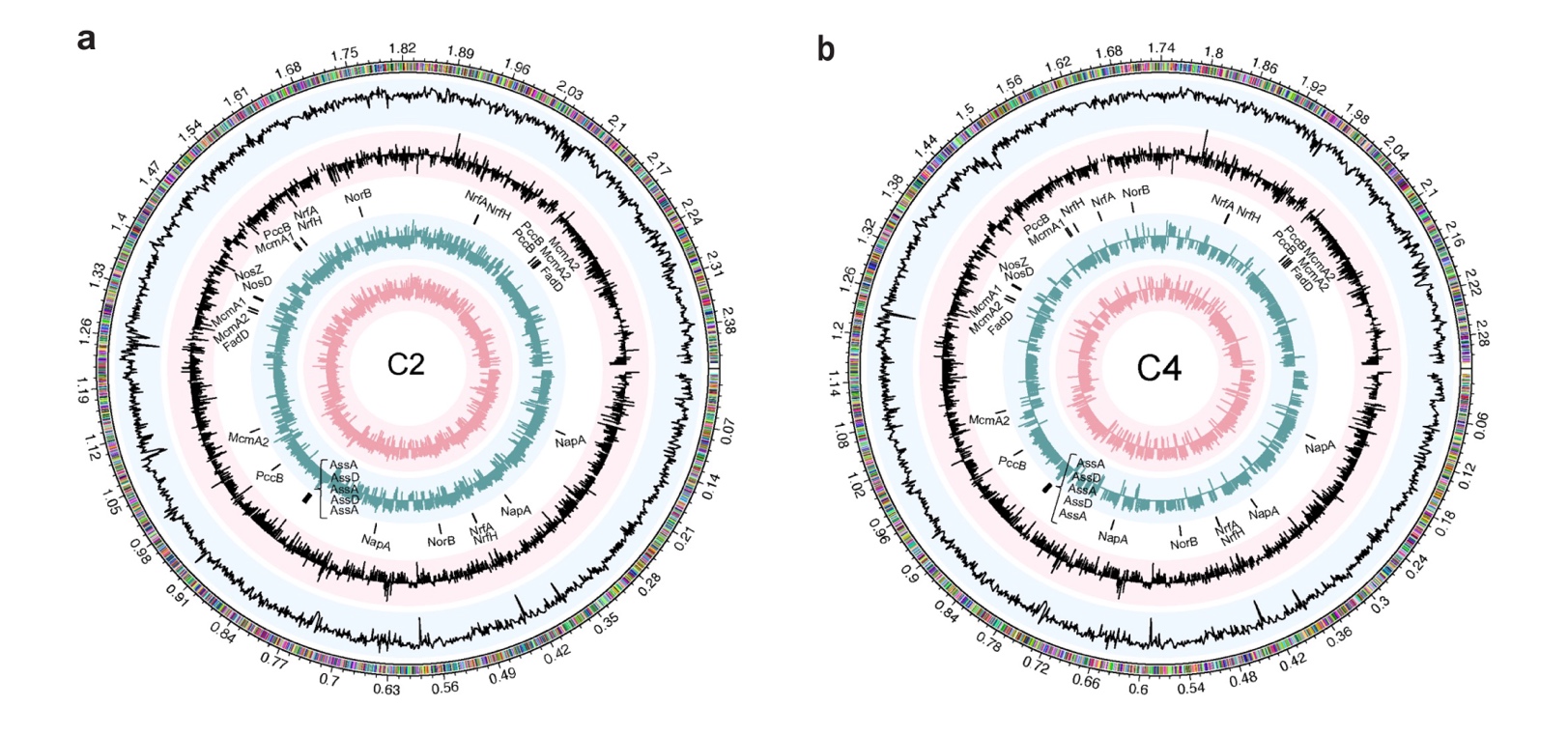

**Supplementary Fig. 6** **Features of recovered Symbiobacteriia genomes in C_2_H_6_- (a) and C_4_H_8_-fed (b) bioreactors.** Tracks from outside to inside: Track 1 - the genomes and encoded genes, Track 2- GC ratio, Track 3 - GC skew, Track 4 - loci of genes involved in nitrogen and ethane/butane conversion, Track 5 and 6 - log-transformed gene TPM values in Phase 1 and 2. The lines in Track 5 and 6 facing outward and inward indicate the genes encoded on ‘+’ and ‘-’ strand, respectively. Ass, Alkylsuccinate synthase; Fad, long-chain acyl-CoA synthetase; Mcm, Methylmalonyl-CoA mutase; Pcc, Propionyl-CoA carboxylase; Nap, Nitrate reductase; Nrf, Cytochrome *c* 552 nitrite reductase; Nor, Nitric oxide reductase; Nos, Nitrous oxide reductase.

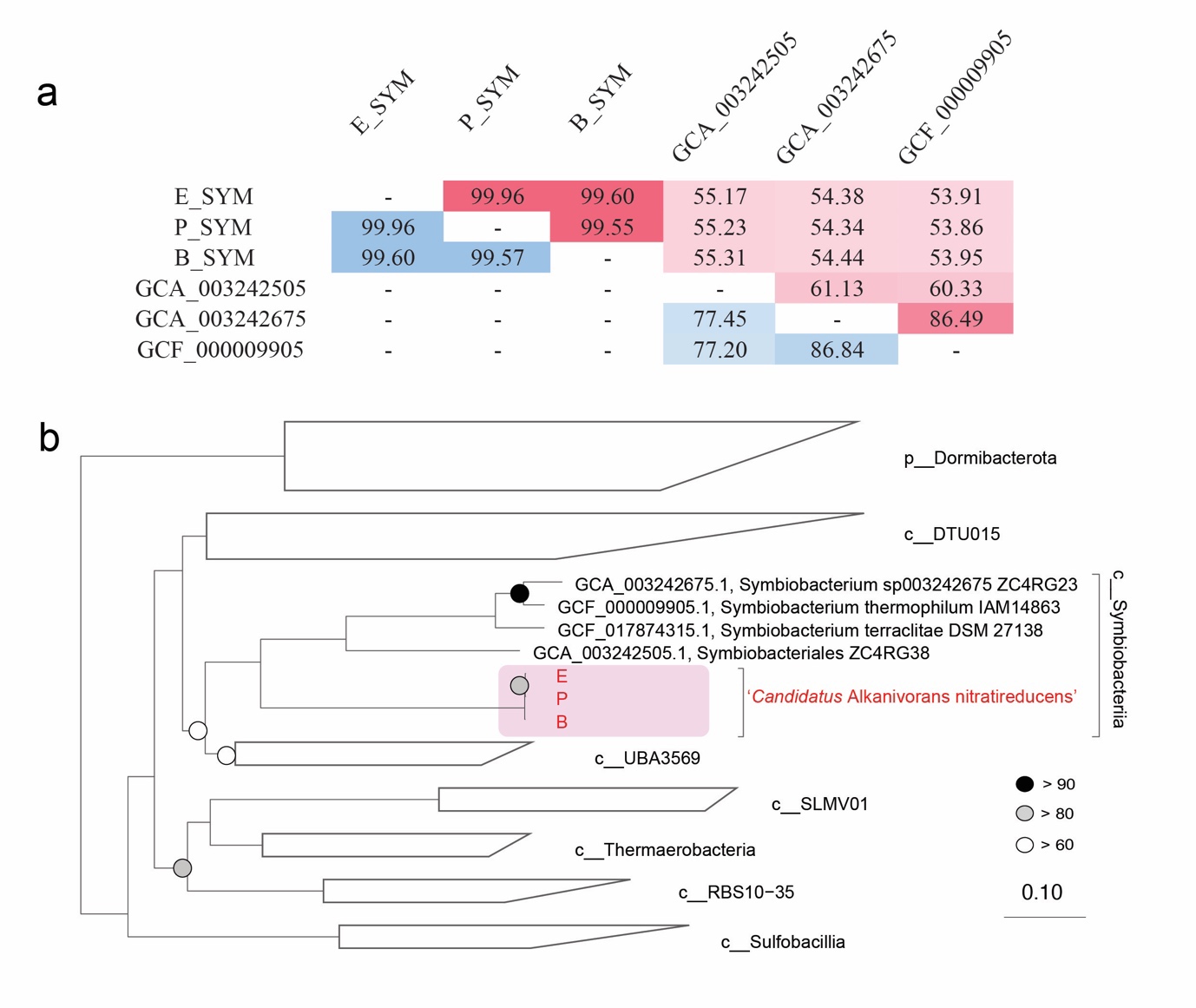

**Supplementary Fig. 7 Comparative genome analyses of ‘*Ca*. A. nitratireducens’ in the C_2_H_6_- and C_4_H_10_-fed systems, and genome-based phylogenetic tree. (a)** ANI and AAI between the available Symbiobacteriia genomes. All available genomes including GCA_003242505.1 (*Symbiobacterium*), GCA_003242675 (*Symbiobacterium*) and GCF_000009905 (*S*. thermophilum IAM 14863) were retrieved from GTDB r202. ANI values were calculated using FastANI, filled in lower matrix and scaled in blue, AAI values were calculated using compareM, filled in upper matrix and scaled in red. **(b)** Genome-based phylogenetic tree inferred with a concatenated set of 120 bacterial-specific marker genes. Genomes recovered from C_2_H_6_-, C_3_H_8_- and C_4_H_10_-fed bioreactors are highlighted in red text. Genomes from phylum Dormibacterota are used as outgroup.

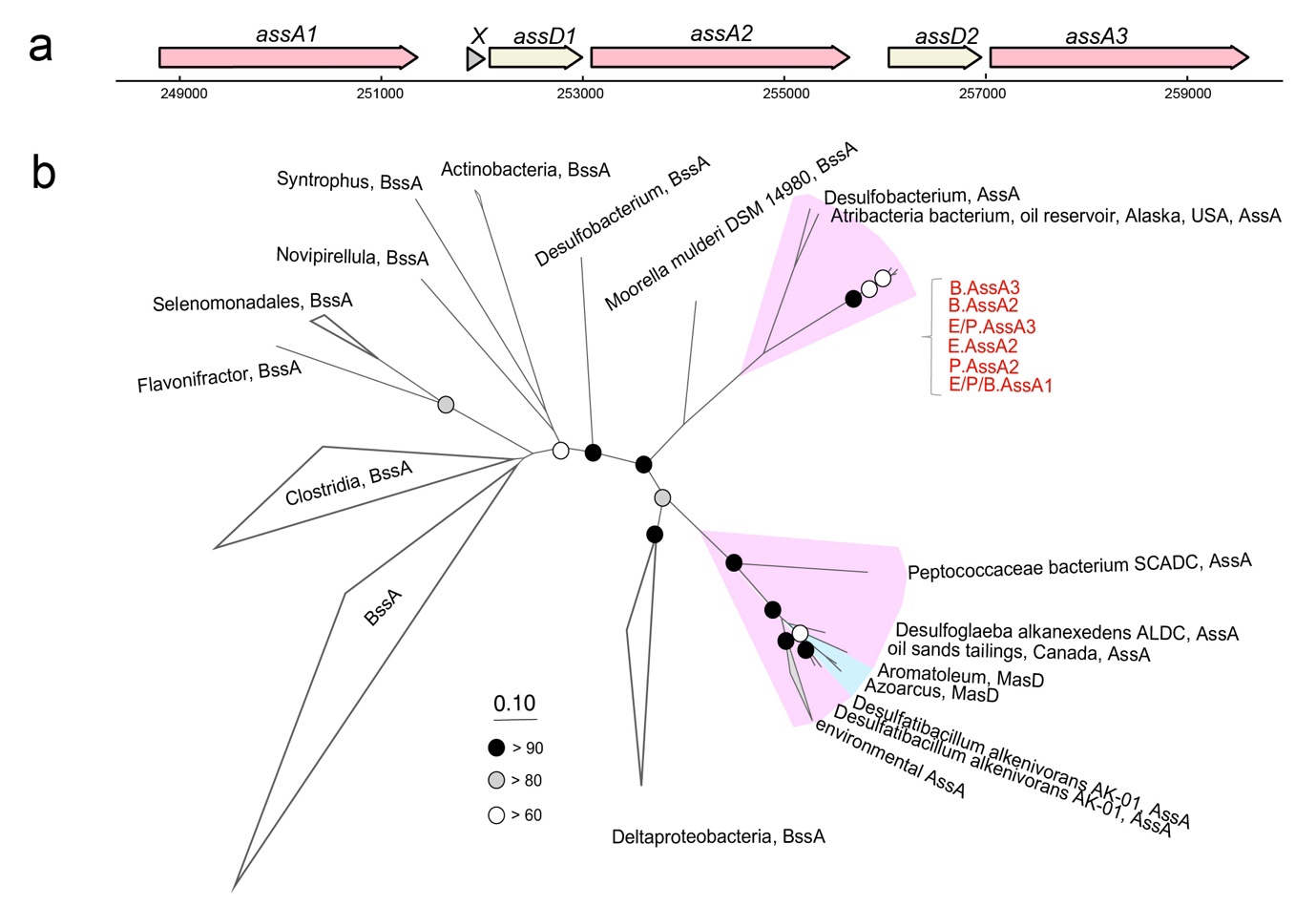

**Supplementary Fig. 8 The operon and phylogenetic affiliation of alkylsuccinate synthase (ASS) in ‘*Ca.* A. nitratireducens’. (a)** The ASS operon in ‘*Ca.* A. nitratireducens’ genome recovered from the C_2_H_6_-fed (E) cultures. Similar with Strain E, the genome of ‘*Ca.* A. nitratireducens’ recovered from the C_4_H_10_-fed cultures also contains three AssA and two AssD subunits. **(b)** Phylogenetic tree of AssA genes. AssA are shaded in pink, while MasD are shaded in blue. AssA recovered from the three Symbiobacteriia genomes are highlighted in red text. Bootstrap values were determined with 100 non-parametric bootstraps, where black, grey and white dots represent > 90, 80-90 and 70-80 bootstrap values, respectively. The scale bar indicates amino acid substitutions per site.

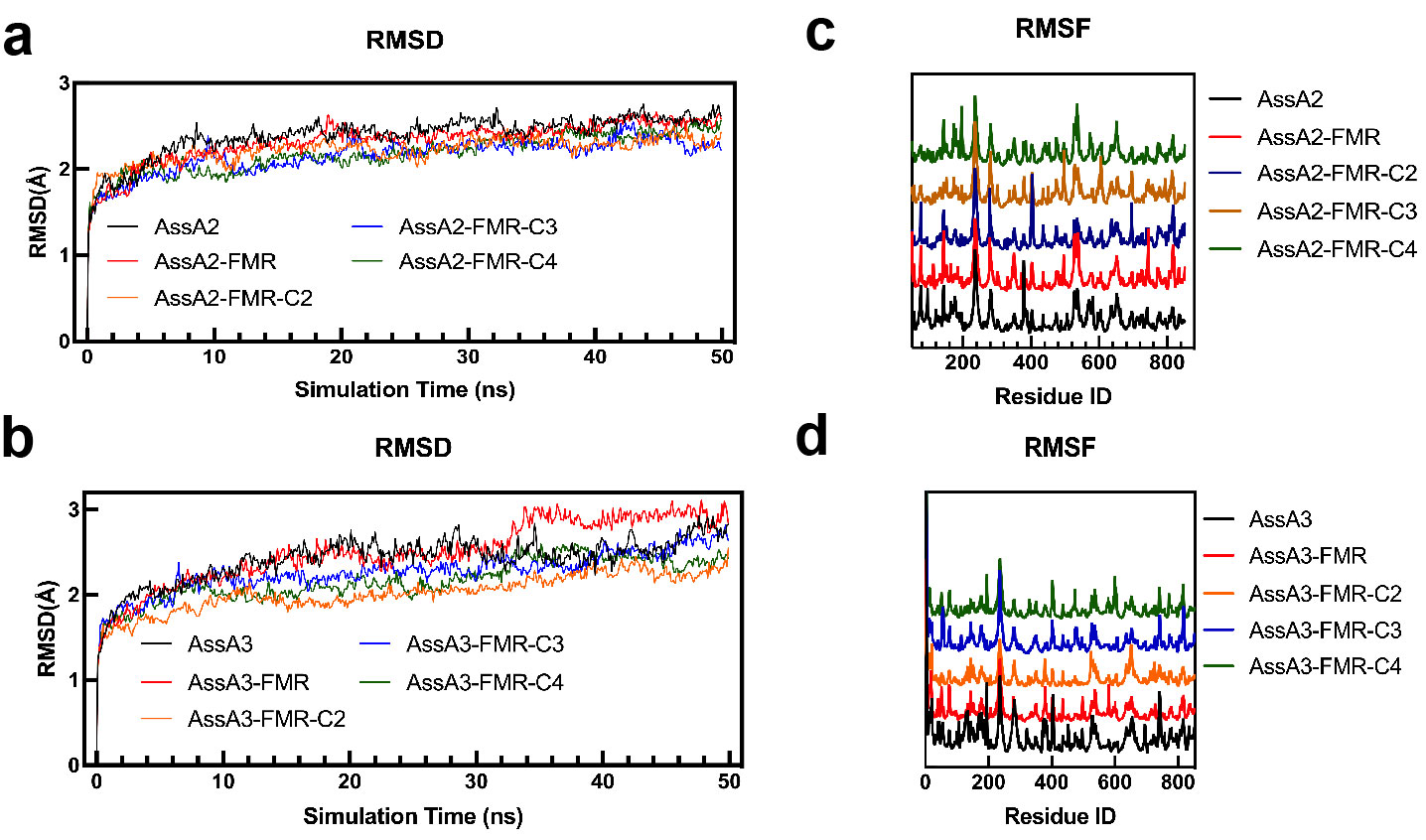

**Supplementary Fig. 9 The calculations of root mean square deviation (RMSD) and fluctuation (RMSF) for all molecular dynamics (MD) simulations. a, b,** RMSD calculations for MD simulations of AssA2 (**a**) and AssA3 (**b**). **c, d,** RMSF calculations for MD simulations of AssA2 (**c**) and AssA3 (**d**). FMR represents the complex with AssA only binding to fumarate, and FMR-C2/C3/C4 means the AssA binds to both fumarate and ethane/propane/butane.

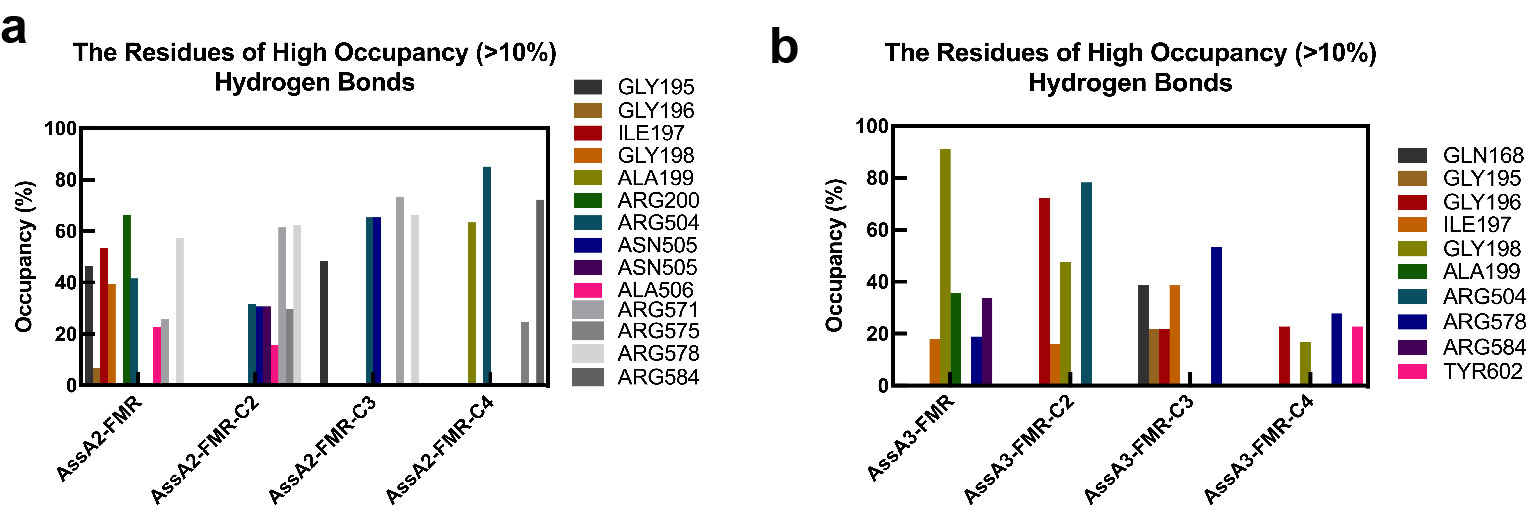

**Supplementary Fig. 10 The amino acid residues with occupancy of hydrogen bonds > 10% in AssA2 (a) and AssA3 (b) in ‘*Ca.* A. nitratireducens’.**

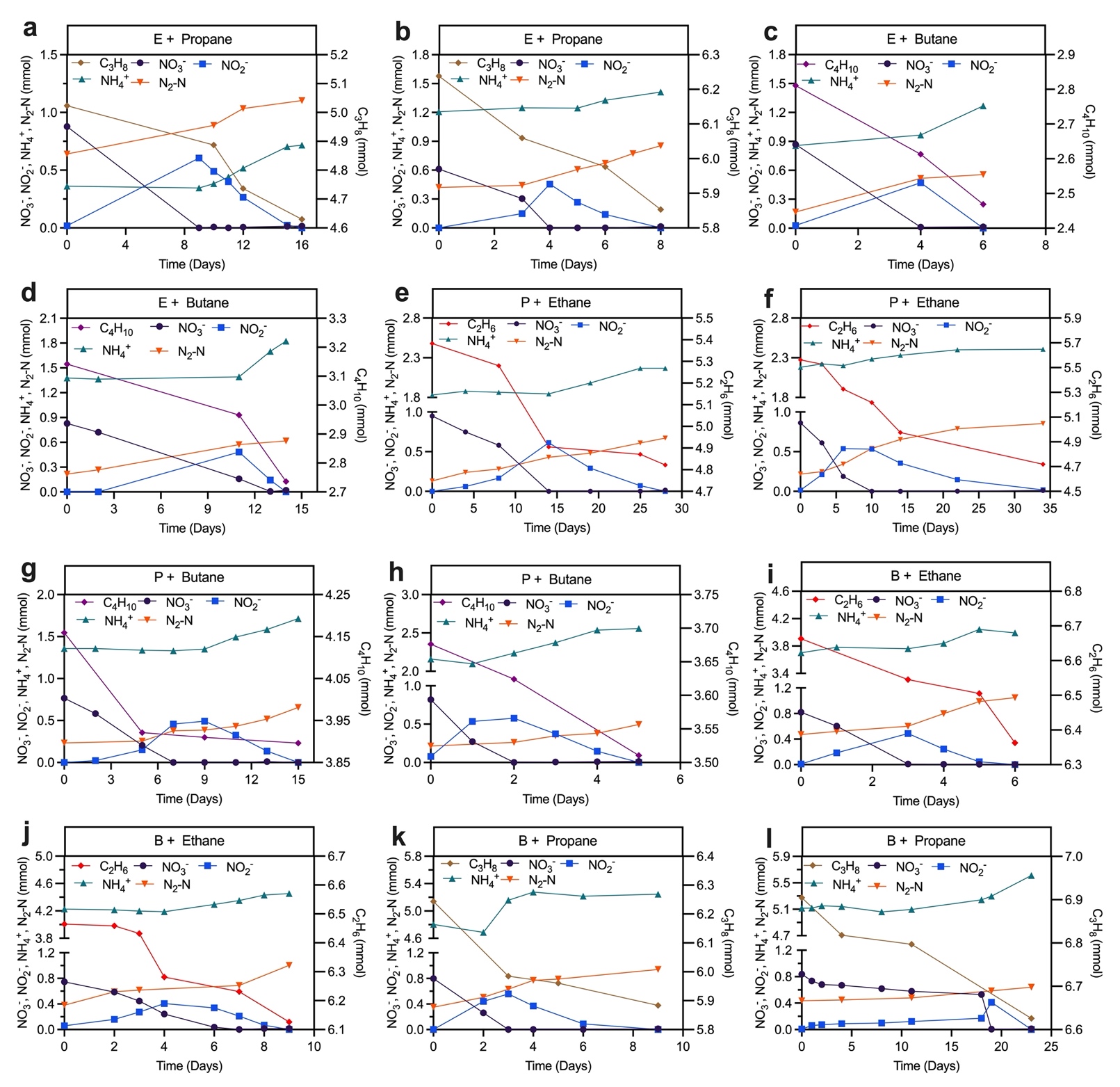
**Supplementary Fig. 11 Profiles of C_2_H_6_, C_3_H_8_, C_4_H_10_, and nitrogen species in the substrate range tests for ‘*Ca.* A. nitratireducens’.**
